# A GPCR negative feedback loop underlies efficient coding of external stimuli

**DOI:** 10.1101/2020.12.14.422627

**Authors:** Rotem Ruach, Shai Yellinek, Eyal Itskovits, Alon Zaslaver

## Abstract

Efficient navigation based on chemical cues is an essential feature shared by all animals. These cues may be encountered in complex spatio-temporal patterns and with orders of magnitude varying intensities. Nevertheless, sensory neurons accurately extract the relevant information from such perplexing signals. Here, we show how a single sensory neuron in *C. elegans* worms can cell-autonomously encode complex stimulus patterns composed of instantaneous sharp changes and of slowly-changing continuous gradients. This encoding relies on a simple negative feedback in the GPCR signaling pathway in which TAX-6/Calcineurin plays a key role in mediating the feedback inhibition. Crucially, this negative feedback pathway supports several important coding features that underlie an efficient navigation strategy, including exact adaptation and adaptation to the magnitude of the gradient’s first derivative. A simple mathematical model accurately captured the fine neural dynamics of both wt and *tax-6* mutant animals, further highlighting how the calcium-dependent activity of TAX-6/Calcineurin dictates GPCR inhibition and response dynamics. As GPCRs are ubiquitously expressed in all sensory neurons, this mechanism may be a universal solution for efficient cell-autonomous coding of external stimuli.

## Introduction

Animals’ fitness critically depends on efficient coding of environmental signals. For example, chemical cues allow animals to locate food sources, a mating partner, and avoid possible dangers. In nature, chemical cues often form complex spatio-temporal patterns. To support efficient navigation based on chemical cues (a process known as chemotaxis), animals need to accurately extract the relevant signals and robustly relay this information to subsequent neural layers.

There are two major modes by which animals may experience chemical cues. In rapid changing environments (e.g., wind carrying plumes or abrupt flow changes), animals will sense instantaneous steep changes in stimulus concentration resembling a step-like function of the stimulus (Murlis et al., 1992). Alternatively, in enclosed and turbulent-free environments, where diffusion processes dominate, smooth chemical gradients will be formed. Under these conditions, animals will typically experience a gradual change in stimulus concentration. In both cases, stimulus concentrations may vary across several orders of magnitude imposing a great challenge for animals to reliably detect and follow stimulus changes(Gaudry et al., 2012; Levy & Bargmann, 2020; Shirley et al., 1987; Sourjik & Wingreen, 2012; van As et al., 1985).

To efficiently chemotax in complex varying environments, living organisms employ at least two sensory-coding principles: exact adaptation and logarithmic coding. Exact adaptation implies that the sensory response to a stimulus is transient, and following an initial change, the response resumes to its baseline levels. This way, the sensory system becomes idle to respond upon encountering impending changes in stimulus levels(Berg & Tedesco, 1975; James E. Ferrell Jr, 2016; Tu & Rappel, 2018; Zufall, 2000). Logarithmic coding, also known as the Weber–Fechner law, is a hallmark of sensory systems found across different organisms and sensory modalities(Fechner, n.d.; Laughlin, 1989). It dictates that perception depends on the ratio between stimulus change and the background level, thus effectively, coding a logarithmic scale of the stimulus. This allows sensory systems to rescale their responses across several orders of magnitude of the signal(Lazova et al., 2011). Interestingly, this feature is thought to be intrinsically implemented in the receptor’s thermodynamic properties, where increased ligand concentrations induce receptor allosteric modulations (e.g. phosphorylation) which shift the binding affinity between the receptor and its ligand(Olsman & Goentoro, 2016).

In single-cell organisms, such as *E. coli* bacteria, these principles are implemented by intracellular signaling pathways to support a biased-random walk chemotaxis strategy(Barkai & Leibler, 1997; Block et al., 1983; Shimizu et al., 2010; Tu, 2013; Tu et al., 2008). In this navigation strategy, cells control the probability for making turns: increasing concentrations of an attractant suppress turning probabilities, such that the cells are more prone to continue moving forward, while decreasing concentrations of the attractant increase turning probabilities(Sourjik & Wingreen, 2012). Crucially, to maintain responsiveness over time, cell activity resumes its basal level even if the stimulus remains constantly on (thus implementing exact adaptation), allowing it to be idle to detect and respond to future changes.

In multicellular organisms, equipped with a neural network, one may assume that similar features may be attributed to dynamics within defined neural circuits. Interestingly however, individual sensory neurons may also implement such computations in a cell-autonomous manner. For example, the *C. elegans* neural network consists of 302 neurons(Cook et al., 2019; White et al., 1986), yet, a single chemosensory neuron, AWA, can cell-autonomously exhibit all these sensory computations to directly control chemotaxis behavior of the animal(C. Bargmann, 2006; C. I. Bargmann et al., 1993; Hart & Chao, 2009; Itskovits et al., 2018; Larsch et al., 2013, 2015).

The AWA neuron shows robust activations in response to diacetyl, a chemoattractant secreted from bacteria in decomposing fruits(Choi et al., 2016). Following a step-like increase in diacetyl concentrations, AWA calcium levels quickly rise, and then return to near-baseline levels even when diacetyl levels remain constantly high (exact adaptation). These responses are found for a wide range of diacetyl concentrations spanning seven orders of magnitude. (Itskovits et al., 2018; Larsch et al., 2015). Interestingly, smooth and slowly increasing gradients of diacetyl lead to a pulsatile activity in the AWA neurons, where the frequency and the amplitude of the pulses increase the greater is the temporal derivative of the stimulus.

As AWA activity facilitates forward locomotion, while a decrease in AWA activity promotes turning events(Itskovits et al., 2018; Larsch et al., 2015), the pulsatile activity dictates a run and tumble strategy, similar to the biased-random walk behavior in *E. coli(Pierce-Shimomura et al., 1999)*. However, while cells typically adapt to the absolute levels of the stimulus, AWA pulsatile activity also adapts to the first derivative of the gradient (Itskovits et al., 2018). This coding principle was shown to support an efficient navigation strategy that outperforms the classical biased-random walk strategy(Itskovits et al., 2018). An intriguing question then arises: how can a single neuron perform all these computations and efficiently encode complex stimulus patterns?

The AWA neuron exclusively expresses the diacetyl G protein-coupled receptor (GPCR), named ODR-10(C. I. Bargmann et al., 1993; Sengupta et al., 1996). ODR-10 activation follows the canonical GPCR signaling pathway: Upon binding of diacetyl, ODR-10 leads to the dissociation of the trimeric G protein, where the Gα subunit stimulates opening of TRPV channels (**Figure 1a**), possibly via polyunsaturated fatty acids (Kahn-Kirby et al., 2004; Larsch et al., 2015). Calcium ions then flow into the cell leading to a partial depolarization that subsequently triggers a much larger influx of calcium through the voltage-gated calcium channels (EGL-19), which culminates in a train of spiking events(Larsch et al., 2015; Liu et al., 2018).

**Figure 1.**
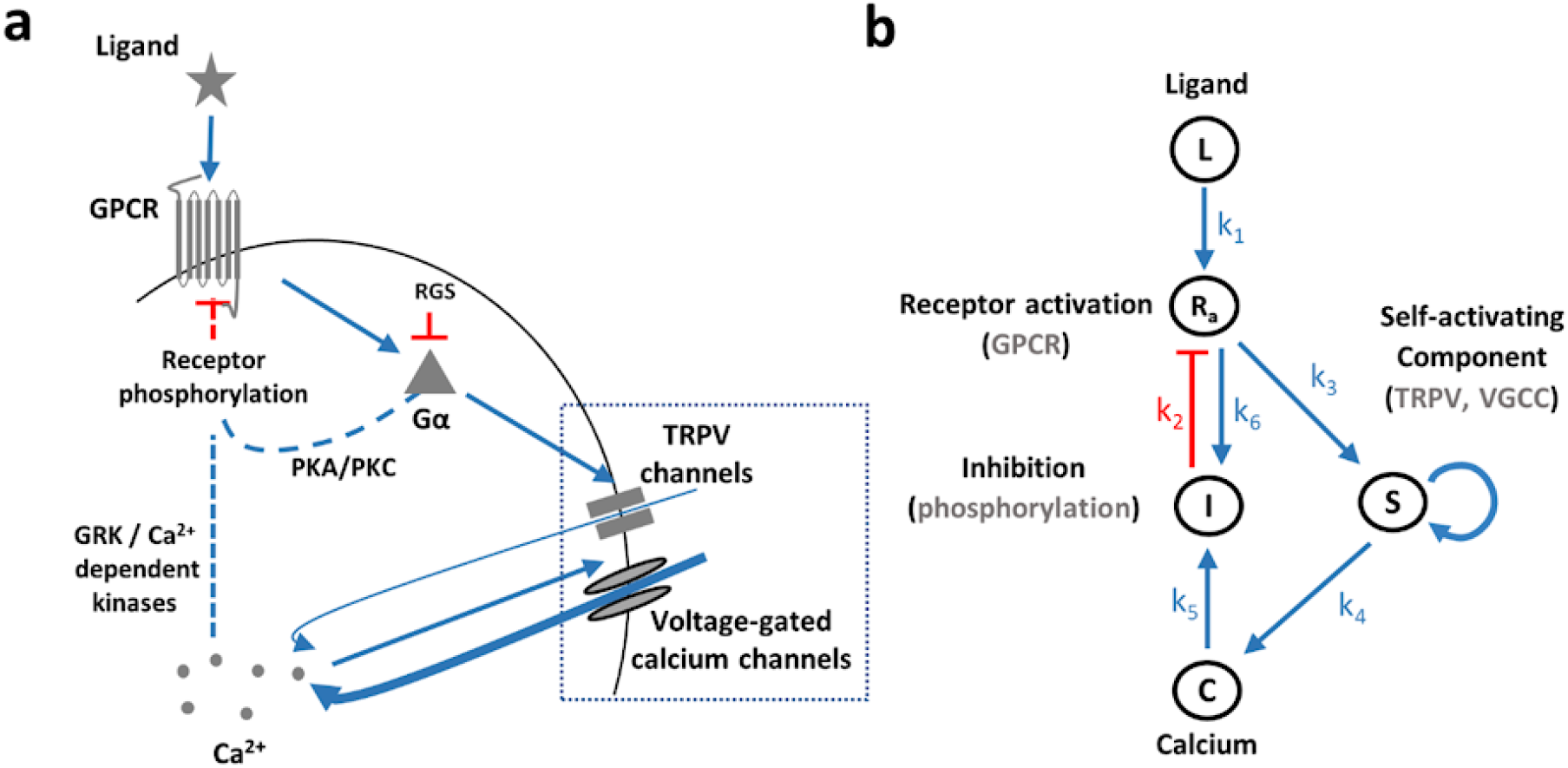
Illustration of the core GPCR signaling pathway and the analogous model. **a.** Key components of the GPCR signaling pathway. Dashed lines mark canonical GPCR pathways, though they have not been verified yet in the AWA neuron. **b.** An analogous circuit topology that recapitulates the main signaling events related to calcium dynamics. Note that S (TRPV and VGCC) depicts a self-amplifying module to generate the pulsatile activity. Calcium-mediated feedback inhibition underlies the exact adaptation. Symbols next to arrows mark the relevant constant in model’s equations. Blue arrows indicate activation and red arrows inhibition.

However, the mechanism by which AWA adapts to the odorant signal is unknown, though evidence suggests that it relies on a negative feedback loop(Rahi et al., 2017). A canonical model for GPCR adaptation relies on receptor phosphorylation events via two possible pathways: (1) second messenger-induced kinases, namely PKA and PKC, which phosphorylate both active and none-active receptors, thus mediating a non-specific (non-homologous) adaptation (**Figure 1a**); (2) G protein-coupled receptor kinases (GRKs) that phosphorylate the active receptors, leading to receptor-specific (homologous) adaptation. GRKs can be activated by calmodulin, a calcium-binding protein, suggesting that calcium levels may indirectly regulate receptor adaptation(Lefkowitz, 1998; Zufall, 2000). In addition, regulators of G protein signaling (RGS) which inactivate the Gα subunit via GTP hydrolysis may also contribute to sensory adaptation(Fukuto et al., 2004; Vries et al., 2000).

In this study, we demonstrate how a simple negative feedback in the GPCR pathway underlies efficient coding of various spatio-temporal patterns of the stimulus. Remarkably, it enables cell-autonomous translation of smooth gradients into a series of pulses that adapt to the magnitude of the gradient’s first derivative. Furthermore, we identified TAX-6/Calcineurin as a key component required for the negative feedback that leads to receptor adaptation. Surprisingly, a simple mathematical model depicts an array of fine neural responses of both wt and *tax-6* mutant animals. Given the ubiquitous expression of GPCRs in sensory neurons, this mechanism may account for efficient coding in other animals across different sensory modalities.

## Results

### A simple feedback model recapitulates all experimental observations for coding complex stimulus patterns

Studies in *C. elegans* worms revealed that a single sensory neuron, AWA, cell-autonomously implements key features required for efficient chemotaxis: (1) It codes the ligand concentration in a logarithmic-like scale, so that neural responses remain similar across orders of magnitude of ligand concentration(Itskovits et al., 2018; Larsch et al., 2015). (2) It responds with a single pulse to a step function and with multiple pulses during a continuous gradual increase in the stimulus concentration (**Figure 2a** and (Itskovits et al., 2018; Larsch et al., 2013, 2015)). (3) It shows an exact adaptation following a step function of the stimulus (**Figure 2a** and (Itskovits et al., 2018; Larsch et al., 2015)). (4) The pulsatile activity (frequency and amplitude) correlates with the first derivative of the gradient and adapts to the magnitude of the first derivative (**Figure 2b** and (Itskovits et al., 2018)).

**Figure 2.**
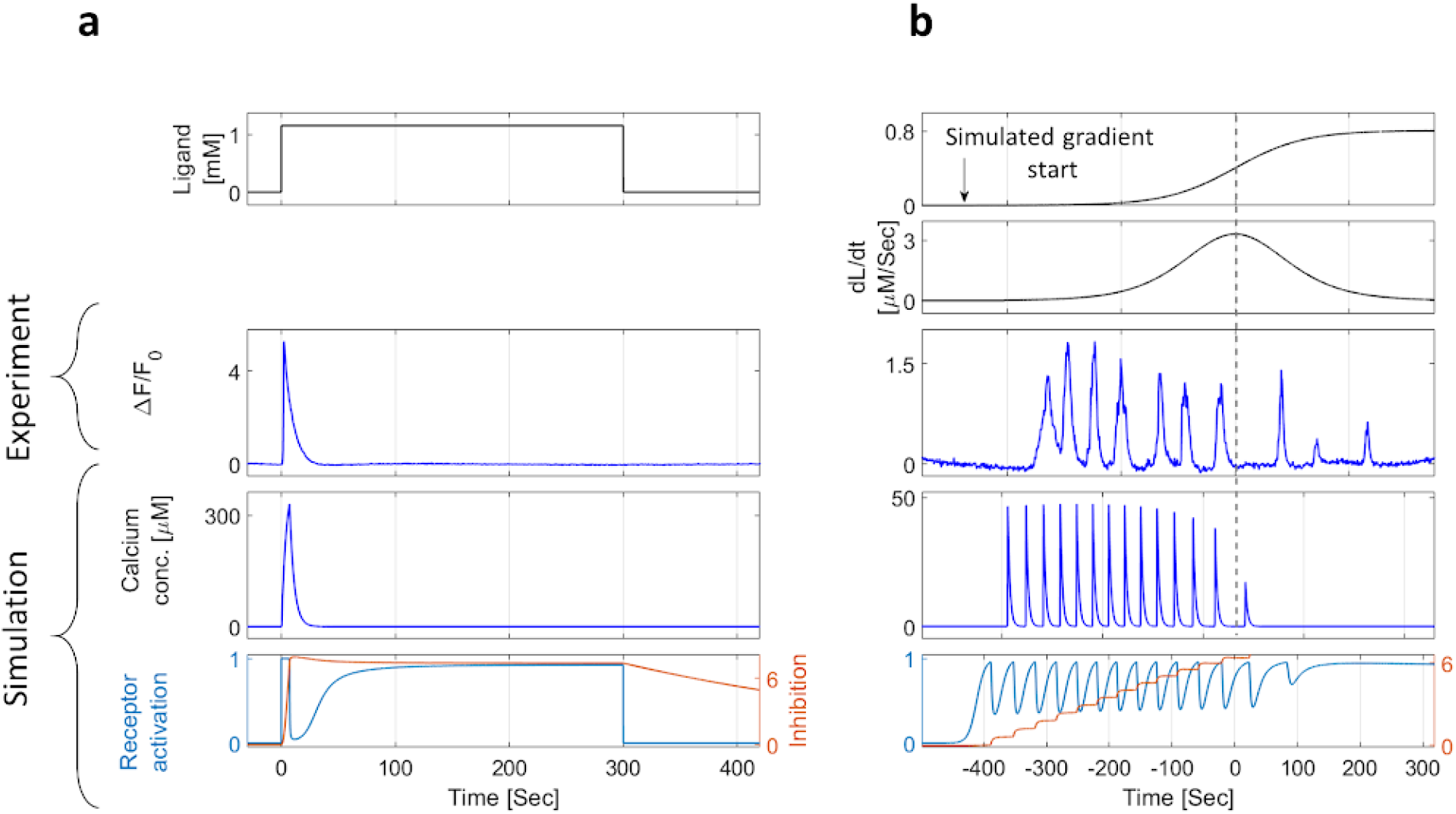
A parsimonious mathematical model recapitulates experimental results demonstrating exact adaptation, pulsatile coding, and adaptation to the magnitude of the first derivative. **(a-b)** Experimental results and model simulations of calcium levels in response to a step function (a) and to a smooth sigmoidal function (b) of the stimulus. Neural activity was measured using a strain expressing GCaMP in the AWA neuron. Diacetyl (stimulus) was presented to the worm using a custom-made microfluidic device (**Methods**). (**a**) Following an on step, and during stimuli presentation, calcium levels rise and then return to their basal levels (exact adaptation in experiment and simulation). Receptor activity (Ra) and inhibition levels (I) reach a new steady state. (**b**) A sigmoidal gradient of the stimulus elicits a series of calcium pulses that are stronger in the first half of the sigmoid, thus demonstrating adaptation to the gradient’s first derivative (dashed line marks the sigmod midpoint, where the first derivative is maximal). This feature is observed in both the experiment and the simulation. The stair-shape inhibition constitutes a discrete memory of previous input levels. Top two panels show the gradient and its derivative. Middle panels depict a representative result of AWA calcium imaging. The two bottom panels present calcium concentrations (C), receptor activation (Ra), and inhibition (I) levels as simulated by the model for the same gradient.

To understand how these coding schemes can be cell-autonomously implemented, we considered the GPCR signaling pathway and constructed a simple parsimonious model of its known signaling components (**Figure 1b**). In this model, the stimulus ligand (L) binds the GPCR to convert it to its active state (R_a_). The active state, through the small Gα subunit, activates TRPV and subsequently VGCC (together denoted as S) to depolarize the cell by increasing cytoplasmic calcium levels (C). Elevated calcium levels activate an array of proteins, including GRKs (denoted as I), which lead to phosphorylation and eventual inhibition of the GPCRs. Gα-mediated activation of PKA and PKC also contributes to GPCR inhibition, and this inhibition is also included in the inhibitory component (I).

The dynamics of this signaling pathway can be simulated using a set of four equations (equations 1-4), where each equation describes the temporal change of one of the main signaling components (see supplementary note for extended descriptions).

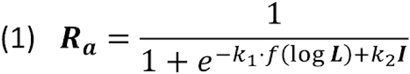

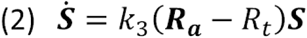

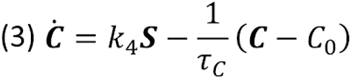

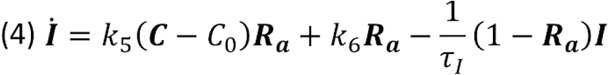

The fraction of active receptors (**R**_**a**_, eq. 1) is described by a sigmoid function and their levels scale logarithmically with the concentration of the ligand (as was shown for the dynamics of *E. coli* receptors(Tu et al., 2008)). **R**_**a**_ inhibition (*e.g.* via phosphorylation) is also linearly dependent on the levels of the inhibitors **I**. This representation is motivated by previous reports showing that when the sensor’s activity increases logarithmically with ligands’ concentration, the negative feedback loop enables logarithmic coding(Tu et al., 2008).

Equation 2 describes a switch-like transition in the state of TRPV and VGCC (**S**). These channels open once active receptors cross a threshold value, R_t_, and close otherwise. The switch-like dynamics is due to self-activating voltage-dependent properties of the VGCCs, through which the majority of the calcium enters the cell. This self amplification underlies the pulsatile activity, and is a known molecular switch motif (J. E. Ferrell Jr & Machleder, 1998; Zhang et al., 2007). The combination of self-activation (Eq. 2) with logarithmic coding (Eq. 1) gives rise to pulsatile activity that logarithmically weakens as the gradient increases, effectively promoting adaptation to the magnitude of the gradient’s derivative.

Intracellular calcium concentrations increase upon opening of the channels (**S**) and decrease once crossing baseline calcium levels C_0_ (Eq. 3). The fact that calcium removal is proportional to calcium concentration enables exponential decay in calcium levels, as was also experimentally observed(Itskovits et al., 2018). Finally, equation 4 describes the circuit negative feedback (**I**), where calcium-dependent (*e.g.,* GRKs) and calcium-independent (*e.g.,* PKA/PKC, denoted by the K6 arrow) pathways enhance inhibition. The full mathematical description and a detailed analysis of the model are available in **supplementary note 1 and supplementary table 1**.

Remarkably, the simple parsimonious model of a negative feedback in GPCR signaling captured all the features that we observed experimentally (**Figure 2**): An on step of the diacetyl stimulus resulted in a single pulse in calcium levels which then decayed to baseline levels despite the fact that the stimulus was constantly present (**Figure 2a**). Similarly, both the experiments and the simulation results showed pulsatile dynamics in response to smooth sigmoid gradients of the stimulus (**Figure 2b**). Moreover, the amplitude and the frequency of the pulses correlated with the gradient’s first derivative and adapted to it. Thus, in response to a sigmoidal gradient, in which the first derivative is symmetric around the gradient’s midpoint, the pulsatile response is stronger in the first half, up to the maximal first derivative point, and decreases thereafter (**Figure 2b, Supplementary figure 1**).

Our model also captures other features observed in AWA response dynamics: In response to repetitive, high-frequency steps of the stimulus, inhibition removal may be too slow and hence activity would not be observed in response to all repetitive stimulations, a phenomena known as periodic skipping (**Supplementary figure 1a-b**). Furthermore, in response to short on-steps, our model predicts gradual habituation (**Supplementary figure 1c**). Both of these phenomena were indeed observed in the AWA neuron, further validating the model (Larsch et al., 2015; Rahi et al., 2017).

The model implements two inhibitory, calcium-dependent and calcium-independent, pathways. The calcium-dependent pathway promotes rapid adaptation that terminates each of the pulses. Blocking this pathway (by setting *k*_5_ = 0) completely abrogated adaptation (**Supplementary figure 2a**). The calcium-independent pathway provides a weaker inhibition through PKA/PKC induced by active receptors. This inhibitory pathway suppresses low-frequency pulses in response to a long-lasting and constant stimulus, as may be seen when setting *k*_4_ = 0 (**Supplementary figure 2b**).

Notably, the model dynamics and its output are robust to changes in the values of the different parameters. We simulated the system behavior while varying each parameter by ~100-fold (See Methods). Despite the broad parameter space, the qualitative dynamics remained largely unaffected, where key features such as exact adaptation and adaptation of the pulsatile activity were maintained (**Supplementary figure 3, Supplementary table 1**). When simultaneously varying all the model’s parameters (except for R_t_) by 10 fold, the qualitative dynamics remained intact in 75% of the cases (see Methods). Moreover, the model’s output is robust to over 10,000-fold variation in stimulus concentration, a necessary requirement for a versatile sensory system that can code and robustly respond to a range of concentrations. These results indicate that the model shows robust outputs that are insensitive to the exact values of its variables.

### TAX-6/Calcineurin is required for the pulsatile response

A classic feedback circuit that achieves full adaptation requires the inhibition to depend on the circuit output(Barkai & Leibler, 1997; Tu et al., 2008). We therefore analyzed several mutants that had been suggested to affect adaptation in *C. elegans* worms. These included: *osm-6*, which is responsible for cilia integrity and intraflagellar transport(Larsch et al., 2015); *eat-16*, an RGS homologue regulating Gα activity; *arr-1*, an arrestin homologue(Fukuto et al., 2004); and *tax-6*, a calcineurin shown to cell-autonomously mediate adaptation in different sensory neurons(Kuhara et al., 2002).

First, we analyzed the capacity of these mutant strains to generate pulsatile activity in response to smooth gradients of the stimulus diacetyl (**Figure 3**). Interestingly, loss of pulsatile activity was observed in *tax-6* mutants only. These mutant animals exhibited a single pulse at the start of the gradient ramp which then slowly decayed over the entire course of the experiment (**Figure 3a-d**). Similar dynamics was observed in some of the *eat-16* and *osm-6* mutants as well, but these responses were not as consistent as in the *tax-6* mutant animals (**Figure 3a,d**). In addition, the maximal amplitude of *tax-6* pulses (as well as of *eat-16* and *osm-6)* was significantly higher than the maximal amplitude observed in wild type (wt) worms (**Figure 3e**). Together, these results suggest that TAX-6/Calcineurin acts to inhibit signaling and reduce intracellular calcium levels, effectively terminating the neural activity pulse. As stimulus levels continue to gradually rise, termination of one pulse supports the generation of the next pulse.

**Figure 3.**
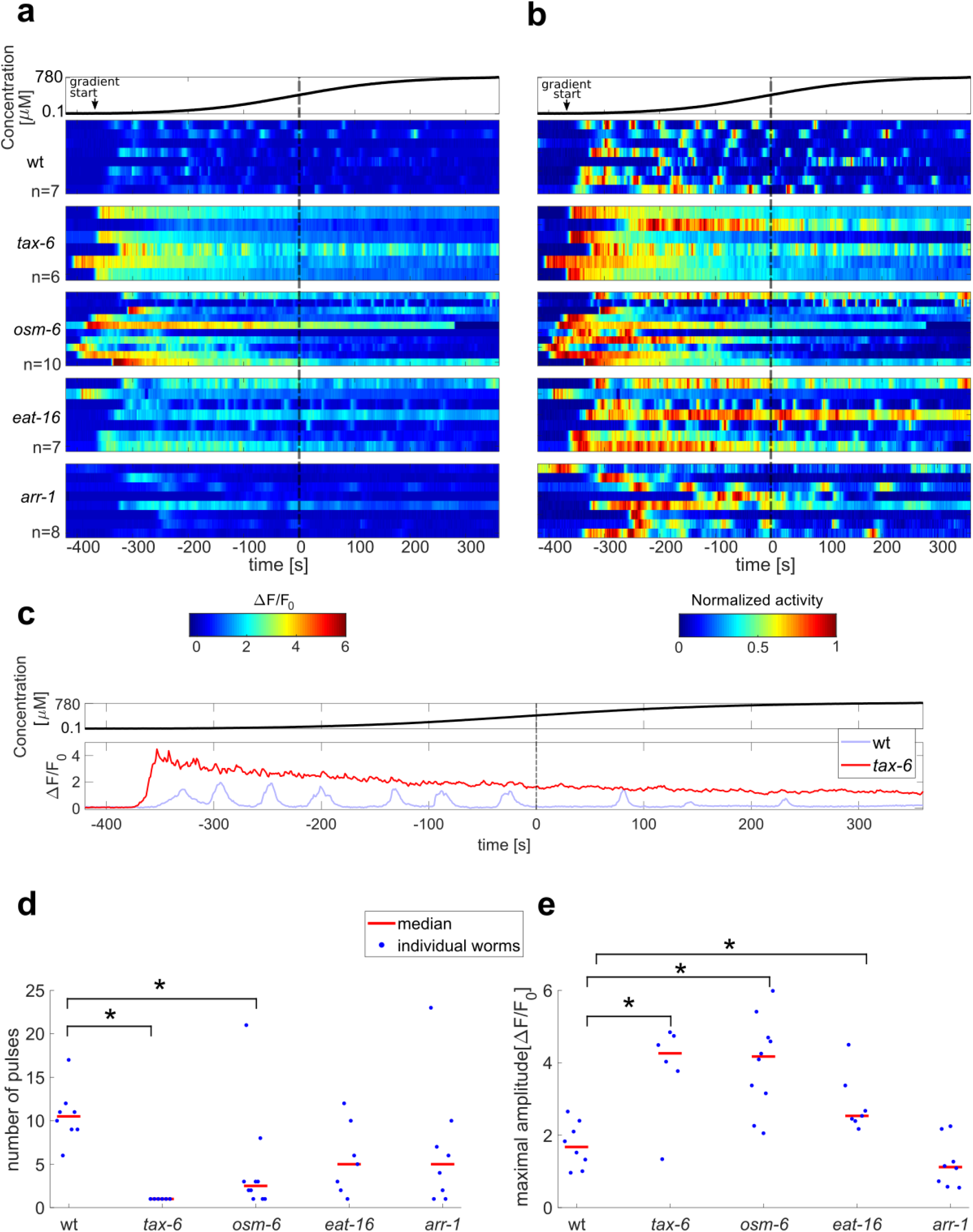
Analysis of dynamic responses and adaptation in different mutant backgrounds. **a-b**. Response profiles of individual wt and mutant worms to a sigmoidal gradient (top). Shown are the fold change activity (**a**) and the normalized activity (**b**) of wt, *tax-6*, *osm-6*, *eat-16* and *arr-1* mutant worms. Notice the lack of pulsatile activity in all *tax-6* mutants. n, number of assayed animals. Time is aligned to the gradient’s inflection point in which the first derivative is maximal (dotted black line). Arrow marks the approximated time in which odorant levels start to rise. (**c**) Representative traces of wt and *tax-6* worms (taken from the first row of each strain in a). **d-e.** To quantify the differences in activity between wt worms and the various mutants, we automatically extracted individual pulses (see methods). Number of pulses was significantly lower in *tax-6* and *osm-6* mutants (p=0.004, 0.01, respectively). Maximal amplitude was significantly higher in *tax-6, osm-6* and *eat-16* mutants (p=0.02, 0.004, 0.02, respectively). Comparisons with wt worms. Wilcoxon rank-sum test, FDR corrected for the 8 comparisons.

### TAX-6/Calcineurin is essential for exact adaptation and habituation

As *tax-6* mutants completely lost ability to generate pulsatile activity in response to smooth gradients, we proceeded with this mutant strain to analyze its capacity to reach exact adaptation, a hallmark of chemosensory coding. For this, we exposed the worms to a five-minutes long on step of diacetyl, after which we turned the diacetyl off for two minutes and then inflicted a second short on step (**Figure 4a**). The first long on-step allowed analyzing the capacity to reach exact adaptation, while the second short on-step, following a two-minutes off step, aimed to analyze possible habituation.

**Figure 4.**
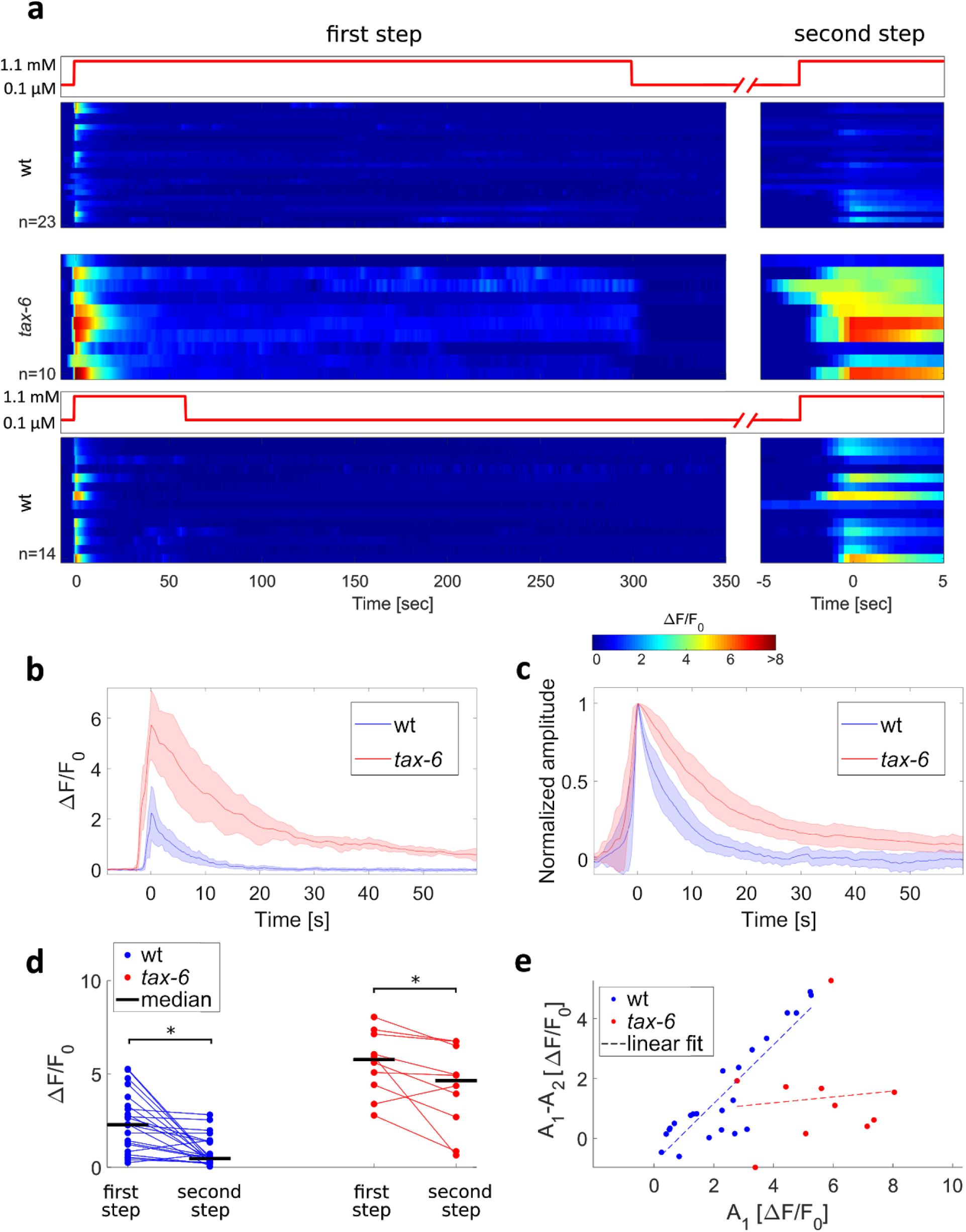
TAX-6/Calcineurin is required for exact adaptation and habituation. (**a**) Response dynamics of wt and *tax-6* mutat worms to a two-step protocol: First step consisting of five minutes (top and middle panels) or one minute (bottom) of on step, followed by a two minutes or six minutes off step, respectively, before applying a second short step (right panels). Activity of wt worms adapted back to its baseline levels in <1 minute following the first five-minutes step, while activity of *tax-6* mutants remained above baseline levels and dropped only when stimulus was removed. Wt worms exposed to the five-minutes step showed higher habituation compared to worms exposed to a one minute step (p=9·10^−4^, Wilcoxon rank-sum test between the amplitudes of the second steps). Steps are aligned to the maximal amplitude (at time 0). (**b-c**) Median neural activity (b) and normalized neural activity(c) of wt and *tax-6* worms in response to the first step. Fitting the pulse with a decaying exponent of the shape *a*·*e*^−*x*/τ^ + *c*, *tax-6* mutants showed a longer decay time than wt worms, suggesting a slower adaptation dynamics (τ _*median*_ of 15 and 5 seconds, respectively, Wilcoxon rank-sum test, p=3·10^−5^). Color-shaded area marks mean absolute deviation. (**d**) Comparison of the amplitudes in the first and the second step for wt and *tax-6* worms (following the longer five-minutes step, top and middle panels in **a**). Both wt and *tax-6* worms showed a weaker activity in the second step (signed-rank test, p=0.00026 and 0.014 respectively), with *tax-6* worms having a significantly higher amplitude in both the first and second steps (Wilcoxon rank-sum test, p=1.8·10^−4^ and 1.1·10^−4^ respectively). **e.** The difference between the amplitudes of the first and second steps is plotted as a function of the first step amplitude. While wt worms show a larger amplitude difference when the response to the first step is stronger (r = 0.89, p=1.3·10^−8^), *tax-6* mutants do not (r=0.1, p=0.78), suggesting that calcium influx affects habituation in wt worms but not in *tax-6* mutants.

Following the first step, the response amplitude of the *tax-6* mutants was significantly higher than that of WT worms, and it was followed by a significantly slower exponential decay **(Figure 4b-c**). Notably, the neural activity of *tax-6* mutants did not return to its baseline levels for the entire duration of stimulus presentation, indicating a failure to perform exact adaptation. Only when removing the stimulus (off-step after five minutes), did the activity resume to its basal level (**Figure 4a**). Importantly, the failure to reach exact adaptation was not due to the enhanced neural activity since *eat-16* mutants also showed enhanced responses to a sigmoidal gradient (**Figure 3c**), and yet, reached exact adaptation following an on-step stimulation (**Supplementary figure 4**).

Furthermore, while both wt worms and *tax-6* mutants showed a weaker response to the second step (**Figure. 4d**), only in wt worms the reduction in response to the second step positively correlated with the magnitude of the first pulse (**Figure 4e**). This suggests that initial stronger pulses, which lead to greater calcium influx, also increase TAX-6-mediated habituation processes **(Figure. 4e and supplementary figure 5)**. Therefore, as TAX-6/Calcineurin activity is calcium dependent, calcium may provide the negative feedback required for the TAX-6-mediated adaptation.

We next asked whether factors other than calcium may also contribute to habituation. For this, we compared neural responses of wt worms following either a short or a long on-step of the stimulus. The short-term stimulus essentially allowed the worms to be off the stimulus for a longer time period before inflicting a second on step (**Figure 4a**). While calcium dynamics in both protocols was similar in response to the first step, the response to the second step was significantly reduced following the long-step protocol where the worms had a shorter time period to recover before the start of the second on-step (**Figure 4a**, **compare top and bottom panels**). Notably, in both protocols, calcium levels between the onset of the first and the second steps were similar. This suggests that a second, calcium-independent adaptation mechanism, may exist. An alternative explanation is that recovery from adaptation can only occur in the absence of an external stimulus. We found similar results when comparing activity dynamics in response to a multi on-step protocol to a single long-step protocol (**Supplementary figure 5**).

### TAX-6/Calcineurin-mediated inhibition does not affect the activity of voltage-gated calcium channels

The response dynamics of *tax-6* mutants to diacetyl was significantly different than that of WT worms: they exhibited a heightened single calcium pulse which did not resume to basal levels (hence no exact adaptation). To elucidate how the lack of TAX-6 inhibition leads to the differential response, we subjected the worms to artificial light-induced activation using the optogenetic channel Chrimson (**Figure 5a**). This light activation increases calcium levels that directly affect VGCC opening, and thus bypassing the natural activation and signaling through the ODR-10 GPCR and the TRPV channels (**Figure 1a**).

**Figure 5.**
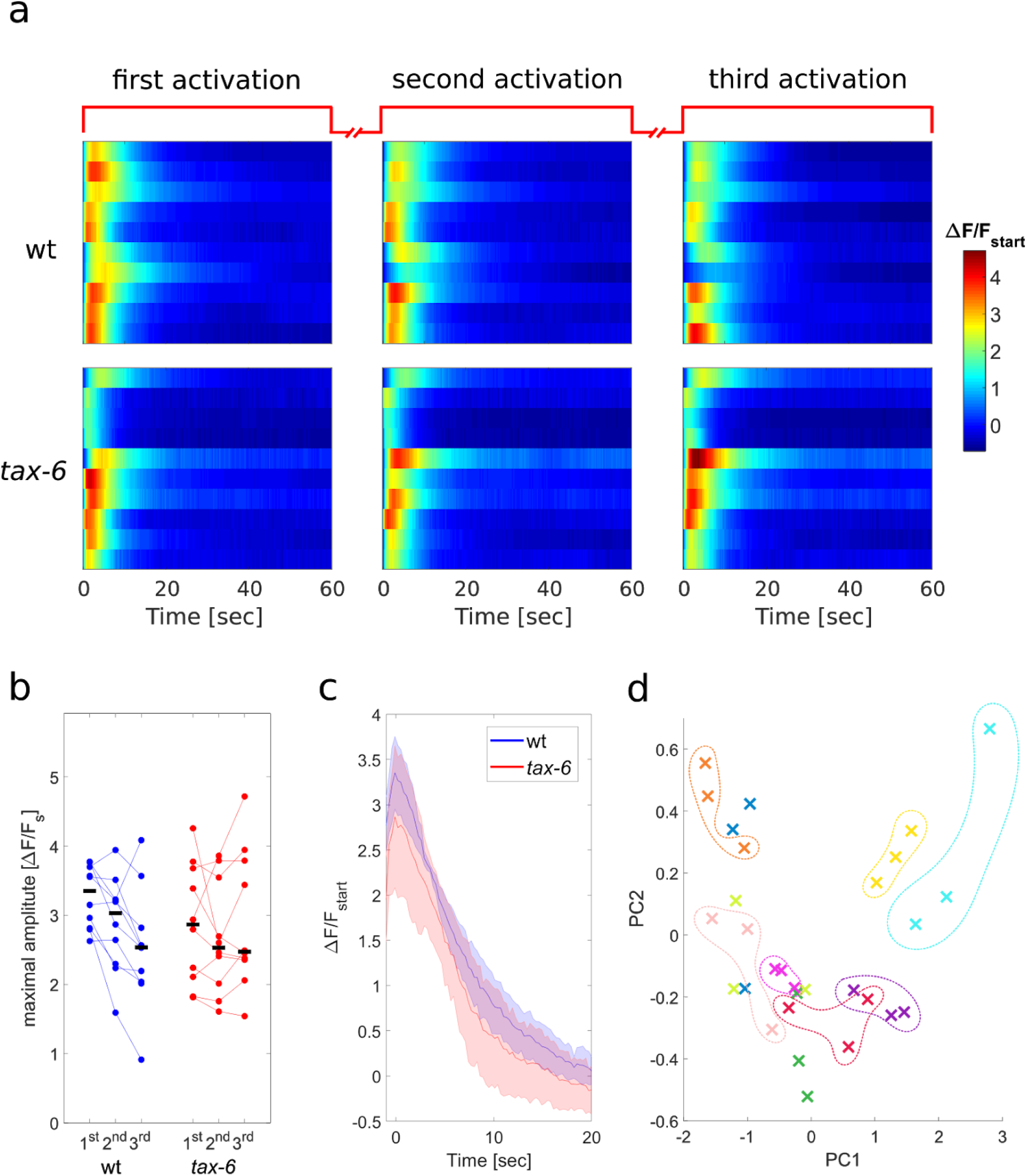
WT and *tax-6* mutants show similar response dynamics following optogenetic stimulation. (**a**) AWA calcium dynamics in response to three repeated light activations (using Chrimson, 488nm). Each worm was illuminated thrice, each for one-minute long with one minute rest between consecutive stimulations. Responses are shown as fold differences in fluorescence from the first captured frame. (**b**) Peak amplitudes as measured for each of the activations. Following light activation, and in contrast to the response to diacetyl, response of *tax-6* mutants was not stronger than the response of wt worms. (**c**) Median activation dynamics of wt and *tax-6* worms. In response to the light activation, the two strains show similar activation dynamics. All responses were aligned to their maximal activation levels prior to extraction of the median values. Shaded-colored area marks mean absolute deviation. (**d**) Activations from the same worm tend to cluster together indicating a low in-worm variability, while the between-worm variability is high. The first 30 seconds in each of the three light activations of the 10 wt worms shown in **a**were normalized by the maximal amplitude and decomposed using Principal Component Analysis (PCA). Shown are the projections of each activation to the first two PCs (capturing 90% and 6% of the variance, respectively). Each color represents a single worm (three activations for each worm). Colored dashed lines were added manually.

For this, neurons were light-activated thrice, each time for one-minute long, while simultaneously imaging calcium dynamics. During the continuous 1-minute long optogenetic activation, both wt and *tax-6* worms showed a similar fast (~2 seconds) increase in calcium levels which was followed by a slow gradual decrease (**Figure 5a**). The second and the third light activations elicited responses that were similar to the responses observed after the first light activation, though with a mildly lower amplitude which may be attributed to bleaching of the fluorescent signal (**Figure 5a-b**). This suggests that calcium influx does not affect optogenetic activation of the neuron, thus placing calcium mediated adaptation somewhere between the ODR-10 receptor and the TRPV channels (**Figure 1**).

Both wt and *tax-6* mutants showed similar neural dynamics in response to light activations (**Figure 5c**), but their responses to the natural stimulus diacetyl significantly differed (**Figure 4b-c**). These findings suggest that TAX-6/Calcineurin mediates calcium feedback inhibition, presumably by affecting the signaling cascade between the ODR-10 receptor (including) and the TRPV channels (see **Figure 1**).

Another interesting observation is the relative heterogeneity in the responses of individual worms to the light stimulus. While the response pattern of each worm seems highly repetitive, the fine activation dynamics in the different worms varied with respect to the amplitude, the rise time, and decay kinetics (**Figure 5a**). To visualize the differences of individual responses, we normalized each response by its maximal amplitude and used Principal Component Analysis (PCA) to project each activation pattern of the WT worms on the two main PCs (**Figure 5d**). Indeed, the three activations of each worm cluster together, indicating a low in-worm variability and high between-worms variability. Notably, these idiosyncratic responses were observed in an isogenic population of worms, expressing the same levels of Chrimson (chromosomally integrated), and grown under the exact same conditions. Thus, the animals’ cell-intrinsic state (*e.g.,* expression levels of the different components, channel’s functional states) governs the shape of the response dynamics.

### A GPCR negative feedback model recapitulates fine dynamic responses of wt and tax-6 mutants

The above experiments position calcium and TAX-6/Calcineurin as key components for GPCR inhibition and exact adaptation. We therefore turned back to our parsimonious model (Equation 1-4 and figures 1-2), and asked whether elimination of TAX-6/Calcineurin from the circuit will recapitulate the experimental results.

To simulate dynamics of *tax-6* mutants, we nullified the calcium-dependent term from equation 4 in the original model (by setting *K*_5_ = 0). In addition, we now included a detailed model for the various voltage-gated channels within the AWA neuron, which for simplicity and analytical tractability, we omitted in the original model (**Figure 1**). This extended dynamics was implemented exactly as it appears in (Liu et al., 2018), who experimentally measured intracellular currents (**Figure 6, supplementary note 1**).

**Figure 6.**
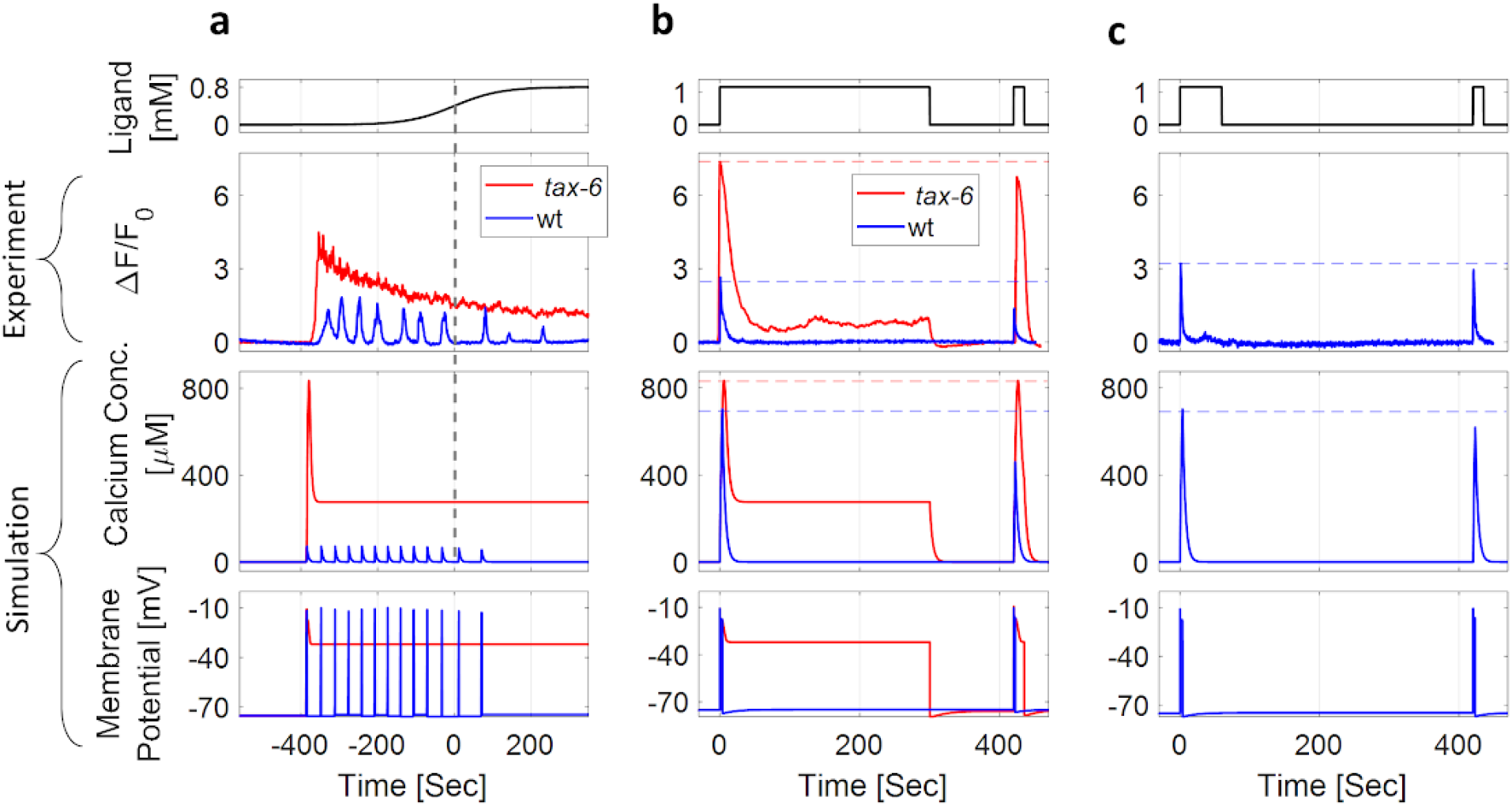
A detailed mathematical model recapitulates experimental results of both wt and *tax-6* mutant worms. In the detailed model (see supplementary note 1), we integrated the available dynamics of the AWA ion channels(Liu et al., 2018). To simulate circuit dynamics of *tax-6* mutants, we set the contribution of calcium mediated inhibition to zero (K_5_ = 0 in equation 4, and shown in figure 1b). **a-c** Calcium concentration and membrane potential as simulated by the model, alongside calcium imaging measurements of a representative worm in response to different stimulation patterns (top). (**a**) In response to a sigmoidal gradient, simulated *tax-6* worms fail to generate pulsatile activity, in agreement with the experimental results. Also, activity does not resume to it’s baseline levels. **b-c**Response dynamics following a long (b) or short (c) two-steps protocol (identical to experiment protocol described in figure 4). (**b**) Experimental and simulated wt worms exhibit lower response (increased habituation) to the second step following the long-step protocol. In contrast, *tax-6* mutants did not show exact adaptation nor reduction in the activity following the second step (no habituation). (**c**) A second on step, following a longer off step period, elicits calcium levels that match the first step. This is in contrast to the short off step period shown in panel b. Despite the fact that in both b and c, calcium levels returned to baseline levels, the longer the off step, the higher the activity response (in agreement with experimental results, figure 4a). This suggests that calcium-independent processes contribute to the inhibition.

The resulting simulations recapitulated the experimental results. When exposed to increasing gradients of the stimulus, WT animals exhibited a pulsatile activity that adapted to the magnitude of the gradient’s first derivative (**Figure 6a**). In contrast, activity of *tax-6* mutants showed a single pulse whose amplitude was higher than that of the wt worms. However, our simplified model failed to accurately describe the slow decay of the pulse in *tax-6* mutants, probably due to our oversimplification of the two-states self-amplification process.

Experimental and modeling results were also in full agreement when considering responses to discrete on-steps of the stimulus (**Figure 6b**). WT animals exhibited a single pulse whose dynamics returned to the basal level (exact adaptation), while *tax-6* mutants responded with a single pulse that was higher in amplitude but which remained above basal level for as long as the stimulus remained on. This heightened amplitude of *tax-6* mutants was also observed following the second step whereas the response of wt worms to the second step was reduced, suggesting that *tax-6* mutants fail to habituate to past-experienced stimuli (**Figure 6c, supplementary figure 6**).

Finally, to verify that our model also captures calcium independent inhibitory processes, we compared the simulated and experimental response dynamics of wt worms in the two consecutive on-step stimulation protocols (short and long intervals, **figure 6c and figure 4a**). While calcium responses to the first on step were identical in both protocols, the amplitude of the second pulse, following a short off-step interval, was lower (**Figure 6c**), in agreement with the experimental results (**Figure 4a**). Notably, the lower amplitude observed following the second step was despite the fact that calcium levels already resumed to baseline levels. This suggests that inhibition is also affected by calcium independent inhibition processes.

Taken together, our parsimonious model accurately captures all of the key features that were experimentally observed: calcium-dependent and calcium-independent inhibition, exact adaptation, habituation, and the corresponding dynamics in *tax-6* mutants which lack inhibitory capacity.

## Discussion

In this work, we demonstrated how a simple negative feedback loop in the GPCR signaling pathway promotes efficient coding of complex stimulus patterns. Moreover, we showed how this coding is achieved cell-autonomously and identified the protein Calcineurin (TAX-6) as key for the negative feedback. A simple mathematical model accurately recapitulated the experimental results observed in both wt and a calcineurin-deficient mutant strain, further underscoring the validity of the model and the interpretation of the experimental results.

Notably, the GPCR negative feedback supports several important computational features, including: (1) Coding ligand concentration on a logarithmic scale, thus enabling adjusted responses across several orders of magnitude of the stimulus. (2) Responding with a single pulse to a step function and with multiple pulses when presented with a continuous increasing gradient of the stimulus. (3) Exhibiting exact adaptation, and (4) the pulstaile activity correlates with and adapts to the magnitude of the gradient’s first derivative. It is therefore remarkable that all these features can be achieved cell-autonomously with a simple negative feedback loop.

These features are embedded in the signaling pathway in a modular fashion (**Figure 7**). Adaptation to the first derivative of the gradient and logarithmic coding are achieved by the inherent activity mode of GPCRs(Olsman & Goentoro, 2016). These receptors are logarithmically facilitated by the ligand, but linearly inhibited by intracellular components. Exact adaptation is achieved by calcium-mediated inhibition. As calcium levels can be viewed as the circuit output, their feedback inhibition underlies the exact adaptation in analogy to the exact adaptation observed in the *E. coli* chemotaxis system(Barkai & Leibler, 1997; Tu et al., 2008). Pulsatile coding of smooth gradients is driven by a self-excitatory element which converts the graded response into a series of pulses. As the negative feedback loop links all these features together, removing a single element, such as TAX-6/Calcineurin, disrupts all these different capabilities.

**Figure 7.**
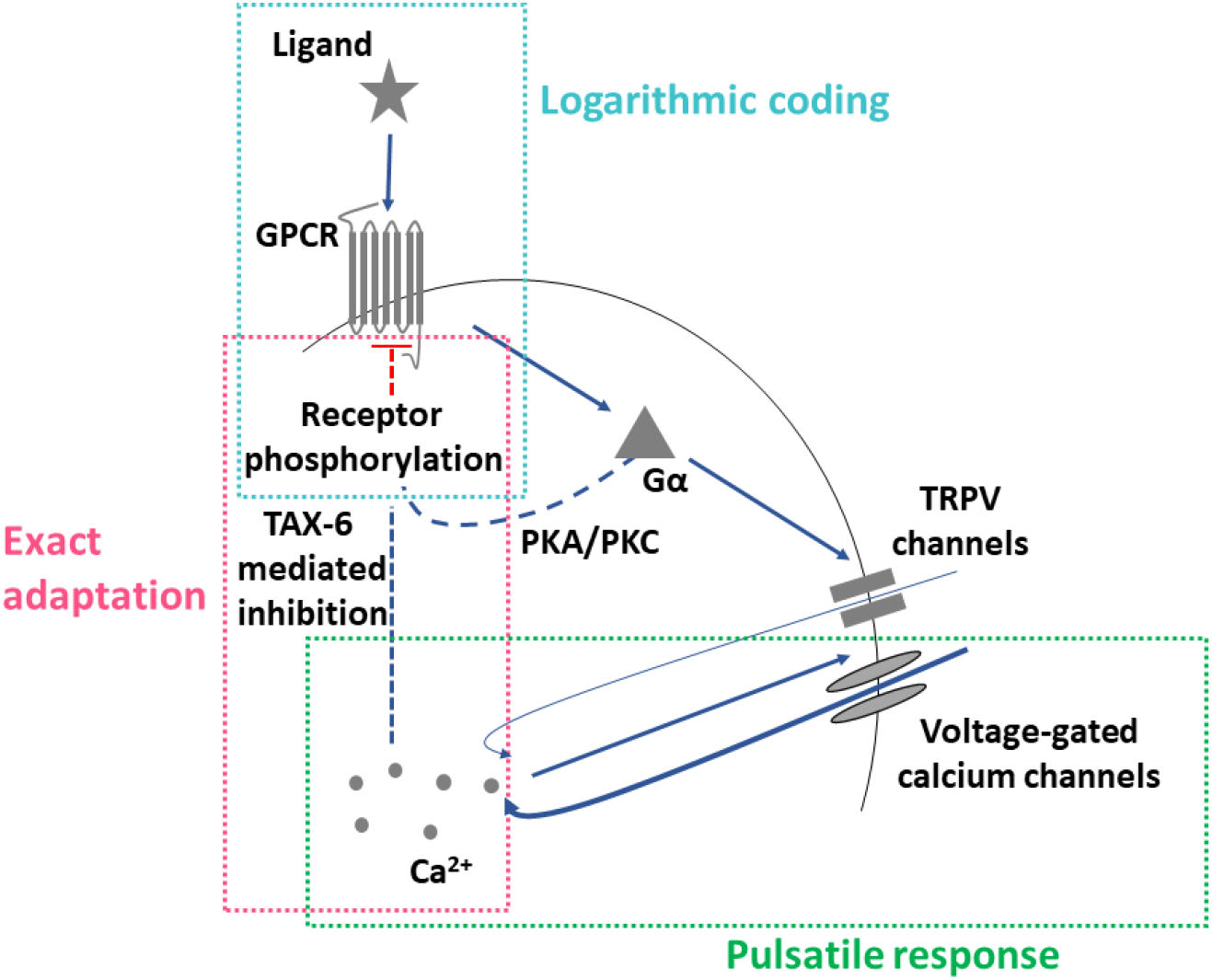
The negative-feedback loop can be decomposed into discrete modules where each module fulfills a defined computational role. TAX-6/Calcineurin and calcium-dependent inhibition mediate exact adaptation. The self-amplifying activity of the TRPV and VGCC channels underlies the pulsatile activity. Logarithmic coding is achieved due to the combination of logarithmic transformation of ligand levels into receptor activity with the receptor’s negative linear dependence on feedback from the system’s output activity (calcium levels).

We discovered that TAX-6/Calcineurin serves as a key component in inhibiting GPCR signaling: while *tax-6* mutants were unable to generate pulsatile activity and reach exact adaptation in response to the natural stimulus diacetyl (**Figures 3-4**), their response to light activation was similar to wt worms (**Figure 5**). This positions the TAX-6/Calcineurin to mediate inhibition of either the GPCR itself, or possibly along the pathway from the GPCR to TRPV activation (included).

How does TAX-6/Calcineurin may promote inhibition? One possibility is by mediating (indirectly) phosphorylation of active GPCRs, a process known as homologous adaptation where the active receptors that initiate the signaling cascade are modified. For example, homologous adaptation was shown in the AWC neuron of *C. elegans* worms, where adaptation to one odorant did not lead to adaptation to a second odorant which is sensed by the same neuron, probably because it signals through a different receptor(Colbert & Bargmann, 1995). In *C. elegans*, GPCR phosphorylation is presumably carried out by *grk-1* and *grk-2* (which may work redundantly as a mutation in *grk-2* alone does not lead to reduced adaptation nor to hypersensitivity(Fukuto et al., 2004)), as well as by the calmodulin-dependent protein kinase II homologue *unc-43(Hobert, 2013)*. Interestingly, phosphorylation of active receptors is thought to underlie the logarithmic coding of GPCRs(Olsman & Goentoro, 2016).

It was previously demonstrated that exact adaptation can be achieved when the output signal directly inhibits the sensor in a negative feedback loop(Barkai & Leibler, 1997; Tu, 2013). As calcium drives the main synaptic output of neurons(Katz & Miledi, 1970; Llinás et al., 1976), it constitutes a preferable candidate to mediate adaptation via a calcium-binding enzyme, such as TAX-6/Calcineurin. The link between calcium and inhibition is further supported by the correlation between calcium influx and the magnitude of the habituation **(Figure 3e)**. Interestingly, a similar correlation between calcium levels and habituation was also observed in mice(Vinograd et al., 2017), suggesting that similar principles underlie these processes in higher organisms. As habituation to the stimulus persisted even after calcium levels returned to their baseline levels (**Figure 4a**), a second, calcium-independent inhibitory process, may also exist.

To better understand the key principles that promote these coding dynamics, we constructed a simple mathematical model that fully captured the dynamics of both wt and *tax-6* mutant worms. Notably, rather than accurately fitting the experimental results, the main purpose of this model was to demonstrate how a simple feedback inhibition in the GPCR pathway can robustly explain an array of sensory responses. The agreement between experiments and simulations demonstrated that calcium-mediated feedback inhibition of the receptor may underlie the exquisite response dynamics: it enables the generation of pulsatile activity as well as adaptation to the first derivative of the gradients; it is responsible for exact adaptation in response to step changes; and it mediates habituation between sequential presentations of the same odorant cue.

When constructing the model, we strived to simplify the detailed signaling cascades, focusing on the key computational components that underlie the observed coding. For this, we grouped components acting together or sequentially and abstracted parts of the system’s dynamics. For example, we modeled VGCCs activity as a self-activating component. A more realistic model would include the positive feedback loop between the VGCCs and the membrane potential (dictated in part by calcium). However, our parsimonious approach sufficed to provide the necessary feedback that generates the observed pulses.

Interestingly, the activation dynamics in individual animals considerably varied, even in response to identical highly-reproducible optogenetic activations (**Figure 5**). This variability is in line with previous reports demonstrating significant differences between individual worms in neural activity patterns and behavioral outputs(Gordus et al., 2015; Itskovits et al., 2018; Luo et al., 2014; Pritz et al., n.d.; Stern et al., 2017). Notably, the assayed worms were isogenic and were grown in the exact same conditions. The fact that these neural responses are cell-autonomou(Itskovits et al., 2018) suggests that a significant variability between individual animals is due to differences in cell-specific internal states(Marder & Goaillard, 2006). Nevertheless, despite the inherent noise and the significant differences in response dynamics, all neurons showed exact adaptation and pulsatile activity. This may suggest cell-autonomous mechanisms that provide robustness in face of noise and variability(Alon et al., 1999). In that respect, the mathematical model suggests that this robustness may be inherently encoded in the negative feedback as the neural outputs were largely insensitive to a wide range of changes in the system’s parameters **(Supplementary figure 2)**.

Together, here we delineated how a negative feedback in GPCR signaling provides an array of key sensory features that together underlie an efficient navigation strategy. We identified TAX-6/Calcineurin as a major inhibitory component and constructed a simple mathematical model that fully depicts the experimental observed results in both wt and *tax-6* mutant animals. While this model simplifies the full dynamics within the neuron, it establishes a convenient framework to understand the principles by which a neuron translates complex stimulus patterns into coherent activity outputs. As GPCR signaling is conserved in all eukaryotes, the same mechanisms may underlie cell-autonomous coding schemes in different sensory modalities across the animal kingdom.

## Materials and Methods

### Strains used in this study

AZS163 [*gpa-6*::GCaMP3, *pha-1*::PHA-1; in a *lite-1;pha-1* background]

The following strains were crossed with AZS163:

PR675 [*tax-6*(p675) IV] yielding the strain AZS282
PR811 [*osm-6*(p811) V] yielding the strain AZS416
JT609 [*eat-16*(sa609) I] yielding the strain AZS403
RB660 [*arr-1*(ok401) X] yielding the strain AZS402

AZ423[*azrIs423*(*odr-7*p::Chrimson::SL2::mCherry; *elt-2*::mCherry); *Ex*(*gpa-6*::GCaMP3)] was generated by first integrating CX16561(*kyEx5662*[*odr-7*p::Chrimson::SL2::mCherry (5 ng/ul, elt-2::mCherry])(Larsch et al., 2015) using UV and backcrossing seven times with WT N2 worms. The resulting strain was then crossed with AZS163. As GCaMP3 signal was weak in worms homozygous for the Chrimson, heterozygous worms were picked for imaging.

AZS424: a cross of AZ423 with PR675[*tax-6*(p675)] which yields the genotype *azrIs423*(*odr-7*p::Chrimson::SL2::mCherry; *elt-2*::mCherry); *Ex*(*gpa-6*::GCaMP3) in *tax-6* mutant background.

Worms were grown at 20 °C on NGM plates that were pre-seeded with overnight culture of OP50 according to (Sulston & Brenner, 1974).

### Preparing reagents for calcium imaging

For calcium imaging of the AWA neurons in response to diacetyl, we used two types of solutions: ‘Stimulus’ and ‘Buffer’. Both solutions were composed of a CTM buffer supplemented with 10 mM Levamisole (Sigma, CAS Number: 16595-80-5). Levamisole was used to minimize worms’ movements in the microfluidic device during imaging. Importantly, Levamisole does not alter AWA response activities as non-sedated worms demonstrate similar pulsatile activity(Itskovits et al., 2018). The ‘Stimulus’ input was also supplemented with 1.15 mM diacetyl and 0.5 μM of rhodamine. The rhodamine dye was used to directly measure the diacetyl gradient experienced by the worms. Importantly, the AWA neuron does not respond to these concentrations of Rhodamine(Itskovits et al., 2018). We also added low levels of diacetyl (1.15 μM) to the ‘Buffer’ solution to reduce the effect of possible abrupt changes in diacetyl concentration that would greatly impact AWA activity.

### Calcium imaging setup

We typically imaged one (either left or right) of the bilateral symmetric AWA neurons using an Olympus IX-83 inverted microscope equipped with a Photometrics sCMOS camera (Prime 95B) and a 40X magnification (0.95 NA) Olympus objective. A dual-band filter (Chroma 59012) and a two-leds illumination source (X-cite, Lumen Dynamics) were used to allow iterative imaging of both green and red channels intermittently. Hardware was controlled using Micro-Manager(Edelstein et al., 2014).

### Generating gradients and steps of the stimulus

To generate smooth gradients of dicetyl, we used two computer-controlled syringe pumps to flow the diacetyl and a diluting buffer into a small-volume (50 μL) mixing chamber(Itskovits et al., 2018). The content of the chamber was stirred by a magnetic bead and its output was flown through the nose of the worm while it was constrained in a custom-made microfluidic device.

Generation of step changes were done in an ‘olfactory chip’(Chronis et al., 2007). Switching between on and off steps was done either manually or using an automated valve (LFYA1218032H, the Lee company) controlled by Arduino.

### Image analysis

AWA activity (green) and rhodamine concentration (red) were each imaged at a rate of 1.4 frames/sec. For step gradients that did not necessitate continuous measurements of diacetyl concentrations, we imaged the green channel at a frame rate of 3.3 frames/sec. Image analysis was done using in-house developed MATLAB scripts. We extracted the fluorescence values of the neuron at each time and calculated the change in fluorescence relative to the fluorescence before the gradient onset (or before the step): (F-F_0_)/F_0_ (as in figures 2-4). Normalization of raster plots to a range of [0-1] was done using [val-min(val)]/[max(val)-min(val)].

### Optogenetic activation

12-24 hours prior to imaging, L4 worms were picked and placed on NGM plates pre-seeded with *E. coli* OP50 supplemented with 100 μM ATR. Optogenetic activation was done via a 40× magnification (0.95 NA) Olympus objective by an X-cite blue led illumination source set to 5% of maximal power. As the same LED light source was also used to image calcium levels, the beginning of neural activation was likely to be observed in the first recorded frame. We therefore quantified neural activity by comparing fluorescent levels to the first frame (F-F_start_)/F_start_.

### Extraction of the pulsatile activity in smooth gradients

Pulses were extracted automatically using the following criterion: a local maximum with an amplitude that exceeds 20% of the maximal measured fluorescence, and which is surrounded by a decay of at least 70% of its amplitude. Data points that obeyed this criterion were considered as pulses. In case no point passed this criterion, the number of pulses was set to 1.

### Parameter’ scan and analysis of the model robustness

To analyze the robustness of our model to variation in the different parameters, we simulated the model’s dynamics using two approaches to scan the parameters space: (i) we systematically varied only one of the parameters in the range of 0.1-10 fold of their initialized value (100 fold-range in total). (ii) In each simulation, we simultaneously varied multiple parameters (except for R_t_, whose dynamic range was found to be relatively small, as seen in **Supplementary figure 3**). Each parameter was drawn independently from a log-uniform distribution, with a range that was set to generate a symmetric 10-fold range around the initially set value (shown in **Table 1**). We used this process to generate 10,000 independent sets of parameters.

For each set of parameters, the model was tested for two features: (1) Exact adaptation in response to a single step (1.15 mM) of the stimulus. Here, the output criterion was to observe a single pulse whose duration was less than 60 seconds. (2) Adaptation of the pulsatile response to the magnitude of the gradient’s first derivative (during a sigmoidal smooth gradient as shown in Figure 1c). The output criterion was to observe anywhere between 3 to 100 pulses, where more than 55% of the pulses initiated prior to reaching the middle point of the sigmoid gradient (the point of maximal first derivative, as expected for adaptation to the gradient’s first derivative).

Pulses were extracted from the simulation output using the following criteria: a local maximum with an amplitude that exceeds 1% of the global maximal amplitude. A pulse’s initiation/termination time was calculated as the time that the output rose/fell more than 10%/90% of its amplitude, respectively. Parameter values that produced activity dynamics obeying these criteria, were considered as valid, thus leading to robust behavioral output of the model.

To analyze the model’s performance in response to various ligand concentrations, we performed each of the above simulations independently while modifying the stimulus amplitude only (the other parameters remained fixed at their initially set values).

## Acknowledgements

We thank Cori Bargmann for providing strains and reagents. Some strains were provided by the CGC, which is funded by NIH Office of Research Infrastructure Programs (P40 OD010440). This work was supported by the Israeli Science Foundation (1300/17), ERCStG (336803), and the Jerusalem Brain Community. AZ also thanks the Greenfield Chair in Neurobiology fund.

---

**Supplementary figure 1.**
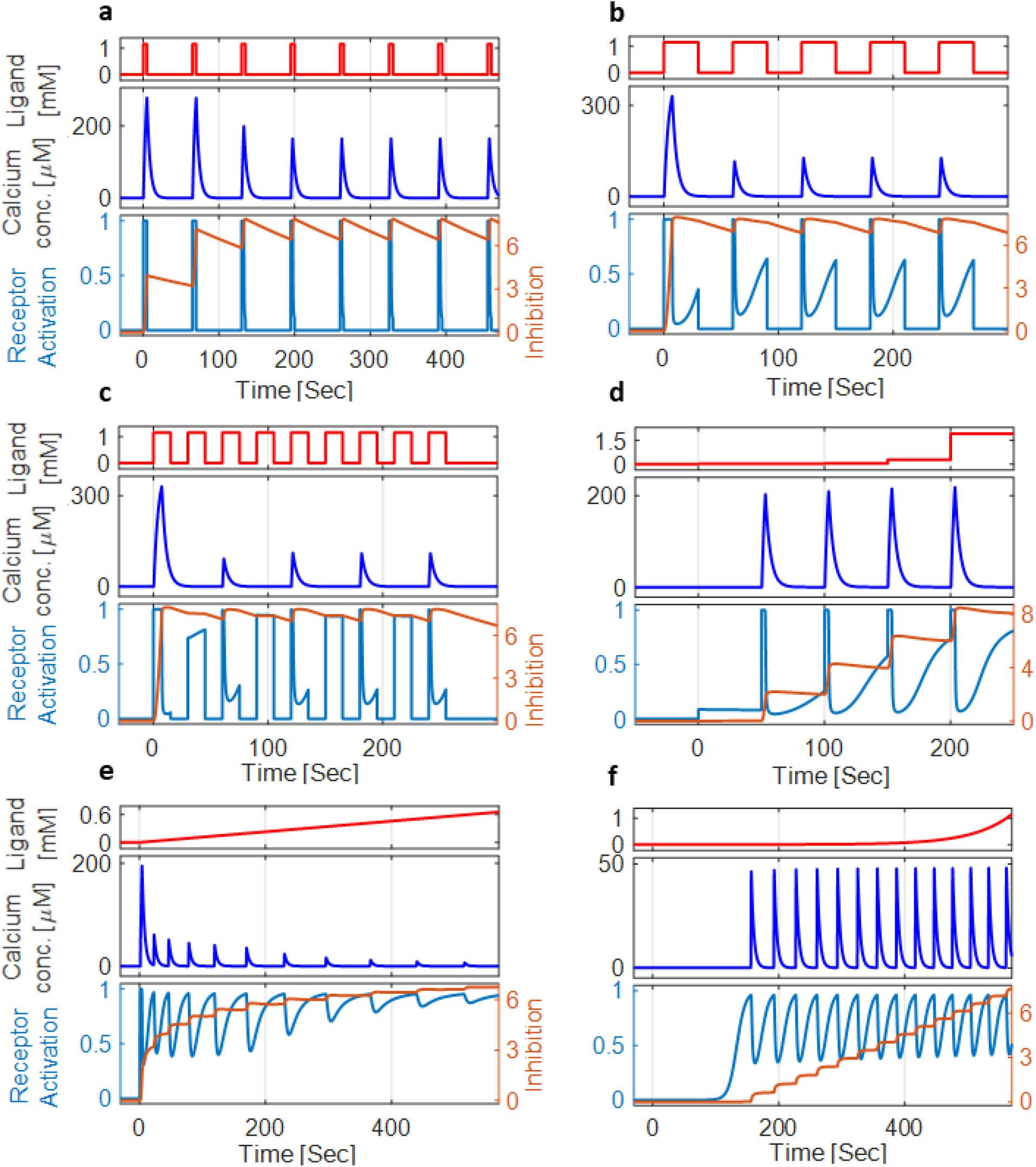
Simulations of the circuit dynamics in response to various stimulation patterns. **a-f.** The circuit output, measured as calcium levels (middle), in response to various input gradients (Ligand, top). Also shown are the simulated levels of inhibition and receptor activation (bottom). **a.** In response to periodic on steps of the ligand (5 s ON and 60 s OFF), the output activity gradually weakens until it reaches a fixed response. The same qualitative response to a similar stimulation pattern was observed in the AWA neuron (Larsch et al. 2015). **b.** When extending the ON-step period and shortening the OFF-step period (30 s ON and 30 s OFF), output activity reaches a fixed response after the first step. **c.** Further shortening the frequency of the stimulus pattern (15 s ON and 15 s OFF) leads to periodic activity skipping, where a calcium pulse is observed in response to every other on-step of the ligand. Skipping occurs because the recovery time from the inhibition (I) during the short OFF period (between two consecutive ON steps) is too short. This is an expected outcome of negative feedback loops that was also observed in the AWA neuron(Rahi et al. 2017). **d.** Nearly-identical output responses when doubling the concentration at each step, reflecting a typical fold-change response. **e.** In response to a linear gradient, the model outcome is a series of pulses whose frequency and amplitude decay over time. **f.** In response to an exponential gradient, neural activity increases with the gradient’s first derivative. Calcium dynamics in e and f is also in agreement with experimental observations(Itskovits, Ruach, and Zaslaver 2018). All ligand patterns start at time 0 and from a baseline concentration of 1.15μM.

**Supplementary figure 2.**
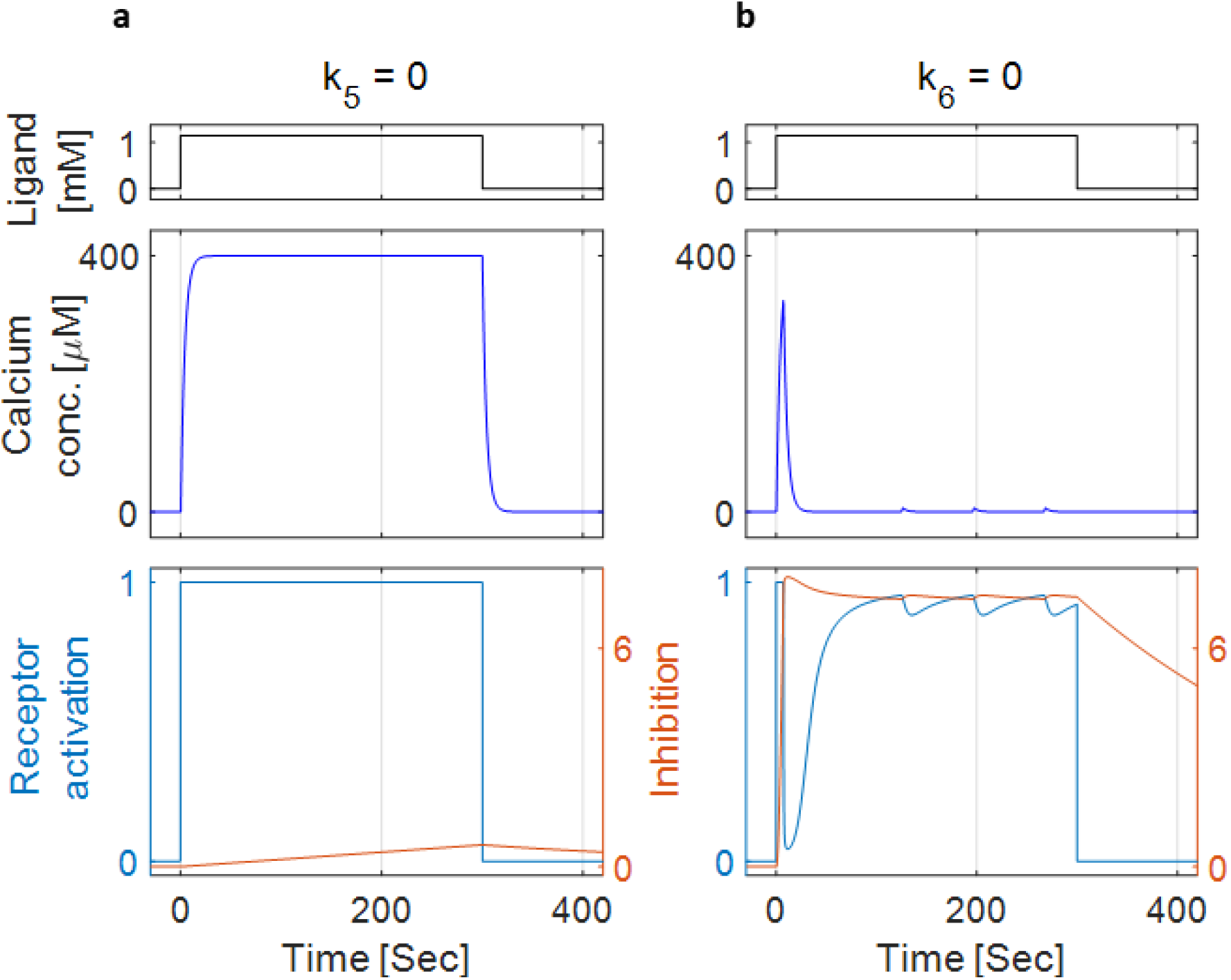
Both calcium-dependent and calcium-independent inhibitions play key roles in the negative inhibition. **a.** When removing calcium-dependent inhibition from the model (setting *k*_5_ = 0), calcium levels do not pulse and remain high for as long as the stimulus is on. **b.** When removing from the model the calcium-independent inhibition pathway (setting *k*_6_ = 0), low-amplitude pulses are observed. Top, a step-function of the stimulus. Middle, simulated calcium concentrations in the cell. Bottom, levels of receptor activation and inhibition.

**Supplementary figure 3.**
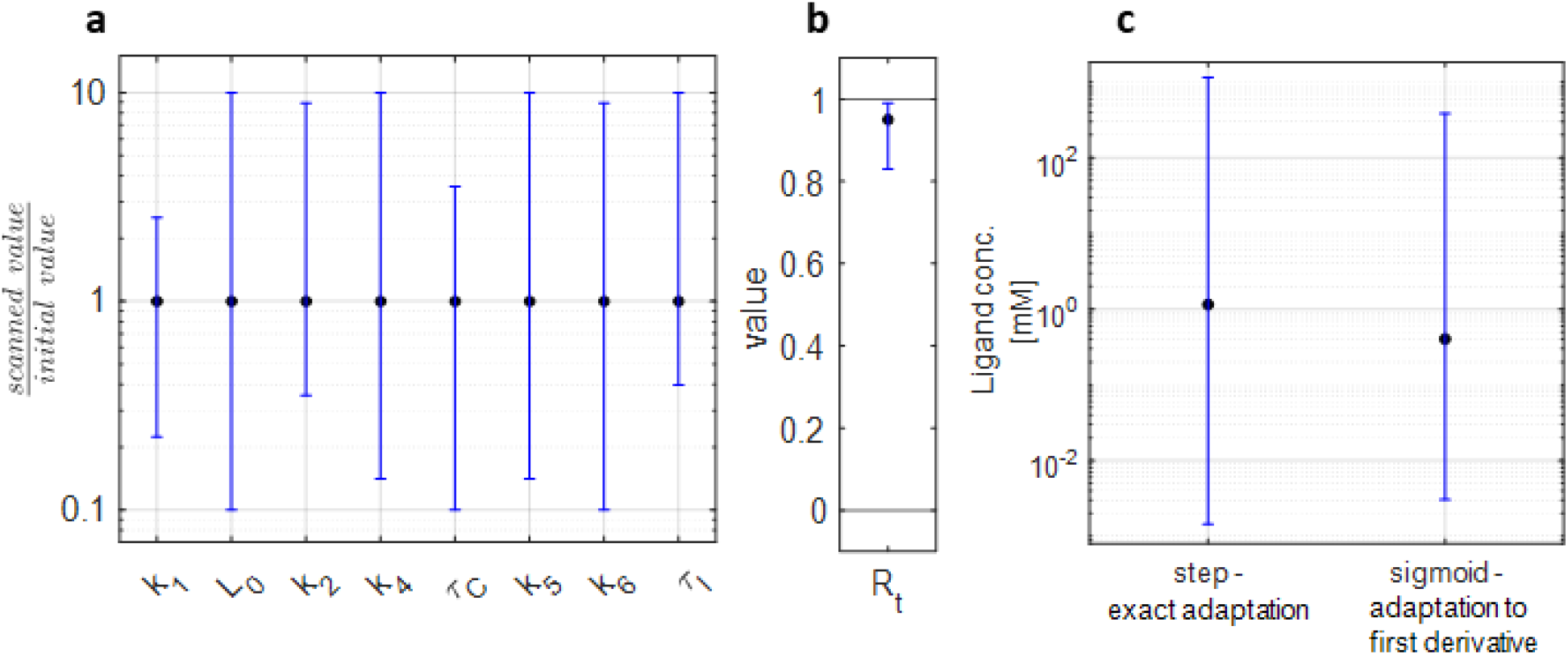
The system’s performance is insensitive to the exact values of the parameters used and performs robustly for a wide range of input concentrations. **a.** Varying each of the model parameters by ~100-fold did not affect the model’s outcome. In each simulation, we varied only one of the parameters, while fixing all the other parameters, and analyzed if the system fulfilled two main features: (1) exact adaptation and (2) adaptation of the pulsatile response to the first derivative of the stimulus (see methods). As the various parameters differed in their units and scales, their ranges are presented as a multiplication of their initial value. **b** *R*_*t*_ values are in the [0, 1] interval and thus shown separately. **c** The range of ligand concentration (almost 6 orders of magnitude) in which the two main features of the model were retained. In all panels, the black dots represent the values used for simulations in this work (presented in table 1) and error bars denote the range of values for which the system fulfilled the above mentioned features.

**Supplementary figure 4.**
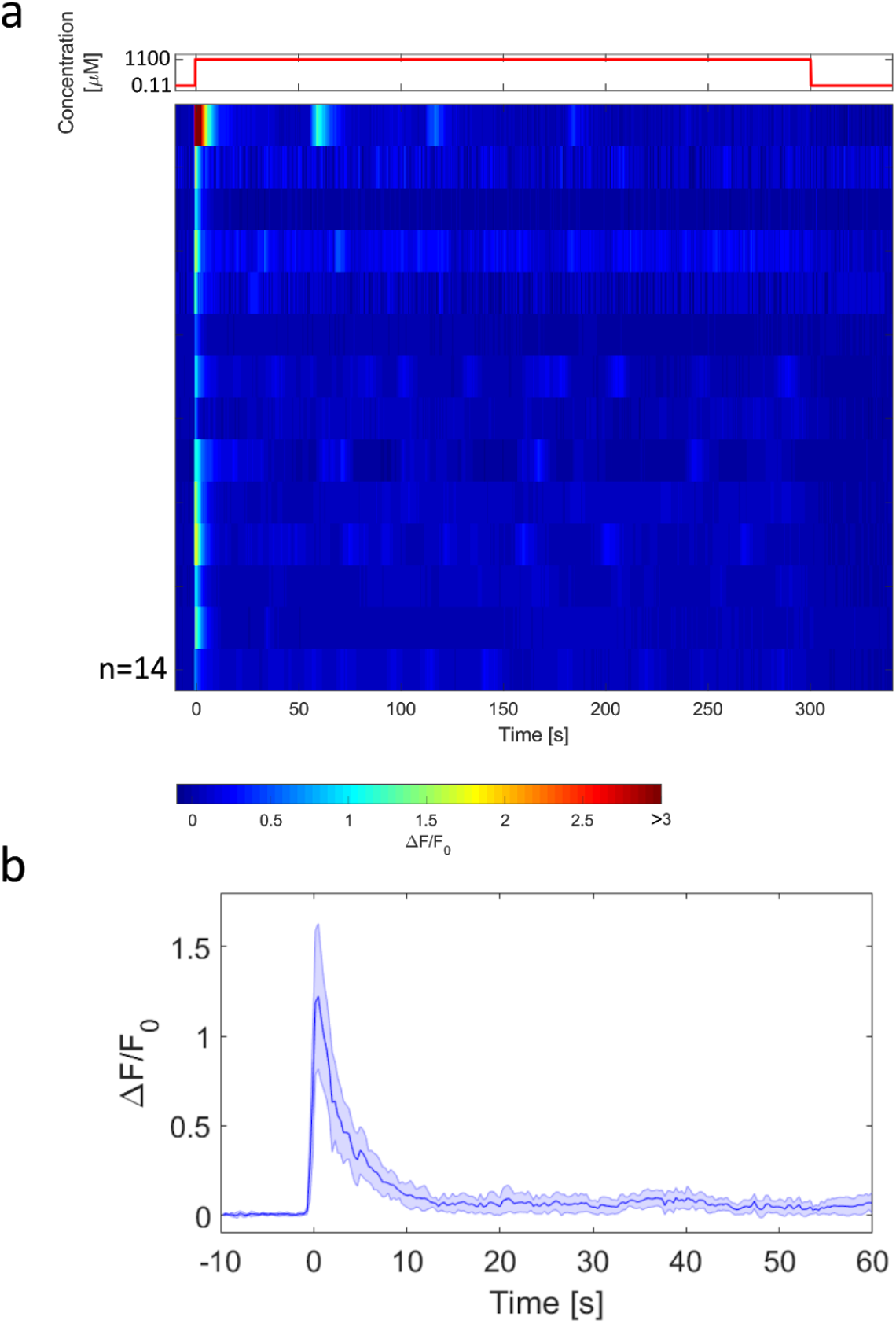
Calcium levels in *eat-16* mutant animals show exact adaptation following an on-step stimulus. **a.** Neural responses of *eat-16* mutants (measured as calcium levels) to a five-minutes step increase of diacetyl (shown on top). While amplitude response in some neurons was low, all animals showed robust increased levels following the ON step. These calcium levels gradually decreased until reaching baseline levels despite the fact that the stimulus remained on. This is in contrast to *tax-6* mutants that fail to return to their baseline activity (as seen in **Figure 4**). The observed low-amplitude pulses may be attributed to the lack of a second, calcium-independent adaptation mechanism (as shown in **Supplementary figure 2**) **b.** Median neural activity of *eat-16* worms in response to the step. Shaded area marks mean absolute deviation. n, number of worms.

**Supplementary figure 5.**
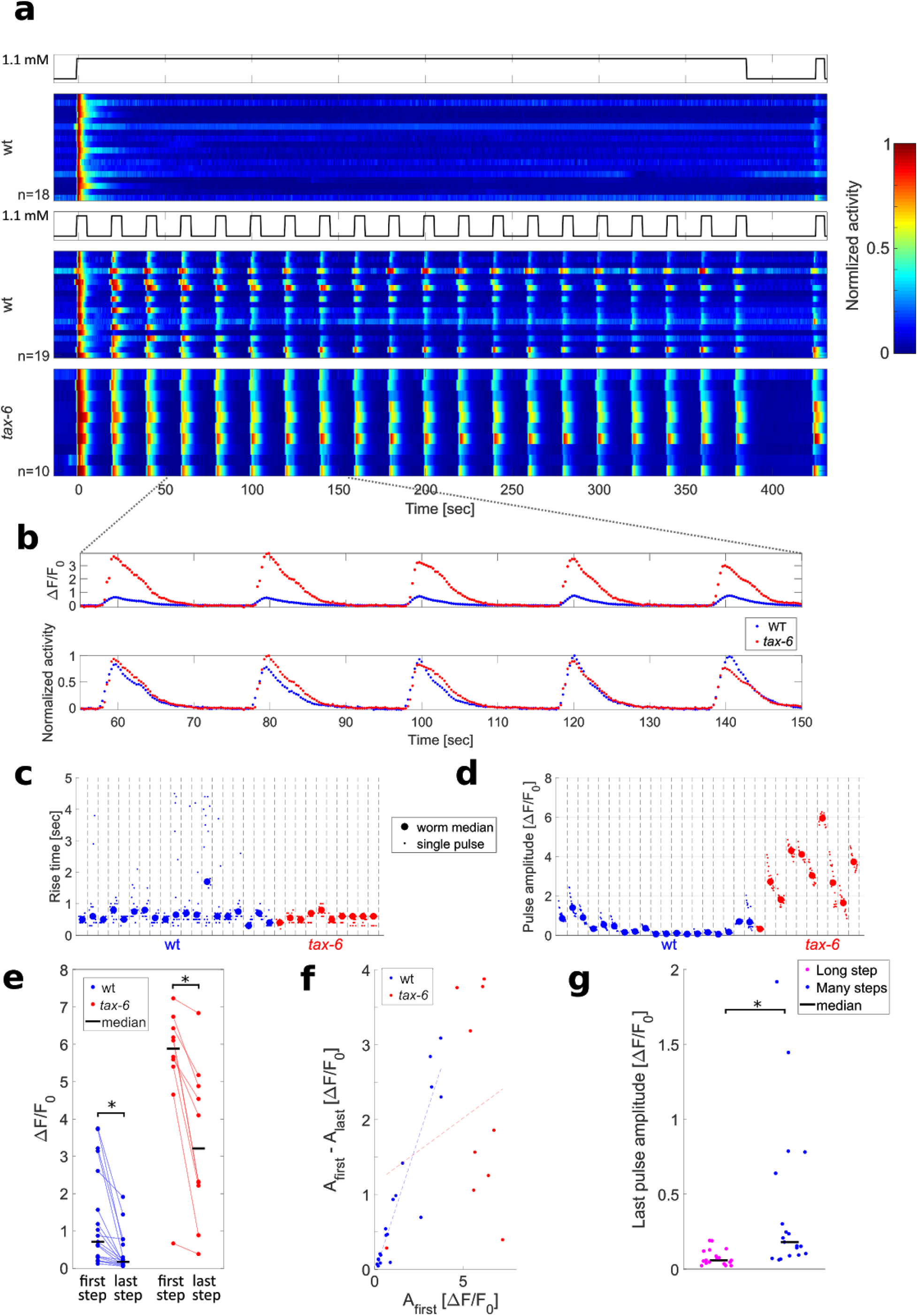
In *tax-6* mutants, the response amplitude is higher than in wt worms, while the response time is the same. **a**. Responses to steps of diacetyl in individual worms. Worms were subjected to either one long step (top panel) or twenty short 5-seconds long steps (middle and bottom panels), followed by a 40-seconds off step and a final 5-seconds on step. Each trace is normalized to its maximal level. **b**. While the pulse amplitude of *tax-6* mutans is higher than that of wt worms, the time to reach the peak is similar. A zoomed-in view of representative responses of wt and *tax-6* worms. Top, absolute values of the response. Bottom, normalized values (by its maximal levels). **c-d**. Quantification of the response time, defined as the time to reach 80% of the maximal amplitude (c) and the peak pulse amplitude (d). While the median response time is not significantly different between *tax-6* mutants and wt worms, the median amplitude of *tax-6* mutants is significantly higher (Wilcoxon rank-sum test, p=9*10^−5^, n = 19, 10 for wt and *tax-6* worms respectively). **e.** Comparison between the amplitudes of wt worms and *tax-6* mutants in the first and last steps of the short interval steps protocol (shown in **a**, middle and bottom panels). In both wt and *tax-6* worms, the response amplitude to the last step was lower (signed-rank test, p=1.3*10-4 and 0.002 respectively), with *tax-6* mutants having a significantly higher amplitude in both the first and last steps (Wilcoxon rank-sum test, p=1.1*10^−4^ and 6*10^−5^ respectively). **f.** The difference between the amplitudes of the first and last steps is plotted as a function of the first step’s amplitude. While wt worms show a larger amplitude difference when the response to the first step is stronger (r = 0.93, p<10^−8^), *tax-6* worms do not (r=0.22, p=0.53). This suggests a link between calcium influx and habituation in wt worms, but not in *tax-6* mutant worms. **g**. Comparison between the neural responses of wt worms in the last step of the long-exposure protocol and the multiple-steps protocol. Mean calcium levels are higher during the multiple-steps protocol (as each step generates a new calcium wave response), and this neural activity is also stronger in the last step following the multiple-steps protocol (Wilcoxon rank-sum test, median of p<2*10^−4^). This suggests that calcium is not the sole factor leading to habituation and additional inhibitory mechanisms may exist.

**Supplementary figure 6.**
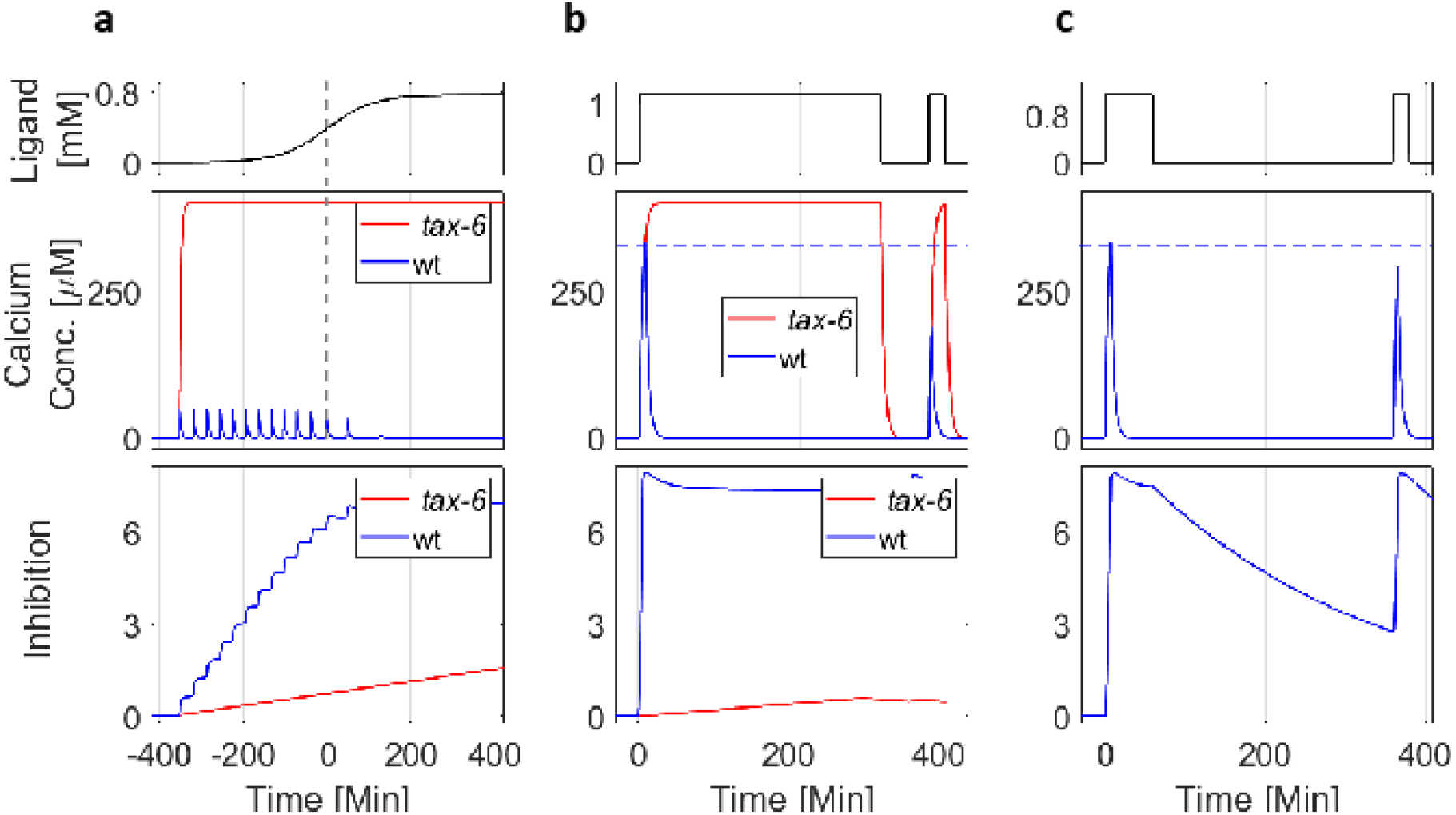
Eliminating the calcium-dependent inhibition from the model recapitulates key responses observed in *tax-6* mutants. **a-c** Simulations of the model outputs (denoted as calcium concentration, middle) and the inhibition dynamics (bottom) in response to various patterns of stimulus presentation (top). To simulate the responses of *tax-6* mutants, we eliminated calcium-dependent inhibition (by setting k_5_ in the model to zero). In contrast to the ‘wt’ dynamics, when removing calcium-dependent inhibition, calcium levels remained at their maximal level for the duration of stimulus presentation, so that exact adaptation was not achieved. **a**. In response to a sigmoidal gradient (top), simulated *tax-6* worms fail to generate a pulsatile activity. **b-c** In a long (b) and short (c) 2-steps protocol, there was no exact adaptation (similar to figure 3 in the main text). Thus, when excluding calcium-dependent inhibition, the simulation’s outcomes match the results obtained using *tax-6* mutants, possibly indicating that TAX-6/Calcineurin mediates calcium-dependent inhibition.

**Supplementary table 1.** The parameters used in the model and their values

| Equation | Description | Parameter | Value [units] |
| --- | --- | --- | --- |
| 1 | Activation facilitation by ligand coefficient | $k_1$ | 25 |
| 1 | Scales the dynamic range of detectable input concentration - determined from(Larsch et al. 2015) by the minimal concentration detected by worms. | $L_0$ | 1 [$\mu M$] |
| 1 | Activation inhibition by modification coefficient | $k_2$ | 10 |
| 2 | Self enhancing component time constant | $k_3$ | 1 [$1/ms$] |
| 2 | Threshold of required Receptor activation to elicit a pulse | $R_t$ | 0.95 |
| 3 | Calcium influx coefficient | $k_4$ | $1 \cdot 10^{-7}$ [$\frac{M}{ms}$] |
| 3 | Calcium removal time constant, determined from(Itskovits, Ruach, and Zaslaver 2018). | $\tau_c$ | $4 \cdot 10^3$ [$ms$] |
| 3 | Steady state calcium level, determined from(Liu et al. 2018). | $C_0$ | 0.1 [$\mu M$] |
| 4 | Calcium mediated inhibition coefficient | $k_5$ | $5 [\frac{1}{M \cdot ms}]$ |
| 4 | Receptor activity mediated inhibition constant | $k_6$ | $2 \cdot 10^{-6}$ [$\frac{1}{ms}$] |
| 4 | Inhibition removal time constant | $\tau_I$ | $3 \cdot 10^5$ [$ms$] |
| 2.1 | Current influx constant for integrated model | $c_{current}$ | 35 [$pA$] |
| 3.1 | Calcium current to concentration change AWA constant (determined from physical properties of the AWA neuron) | $c_{AWA}$ | $2.6 \cdot 10^{-8}$ [$\frac{M}{ms \cdot pA}$] |

## Supplementary note 1. A detailed description and analysis of the mathematical model

Our model for chemo-sensation in the AWA neurons is composed of a system of time dependent differential equations. The circuit’s topology (**Figure 1**) is analogous to the negative feedback loop circuit found in the chemosensory system of *E. coli (Tu, Shimizu, and Berg 2008)*. We therefore capitalized on the mathematical framework developed in these studies and modified it to adjust to the known GPCR signaling pathway found in the AWA neurons of *C. elegans* worms.

Our model consists of five variables. The “Ligand” (L) denotes the input stimulus (*e.g.,* diacetyl). The Receptor Activation term (R_a_) represents the fraction of active ligand-bound receptors in the cell membrane. R_a_ is facilitated by ligand binding and is regulated by Inhibition (I) which represents the effect of inhibiting factors. Overall, R_a_ is modeled as a sigmoidal function of the difference between the Ligand and Inhibition terms:

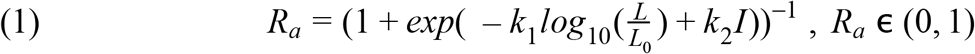

Where *k*_1_, *L*_0_, *k*_2_ are positive constants. In this functional form, *I* serves as the system’s “memory” of past ligand concentrations. The logarithmic-scale coding of the ligand and the linear regulation of the negative feedback (“Inhibition” motif) are concepts borrowed from chemosensation models in *E. coli(Shimizu, Tu, and Berg 2010; Yi et al. 2000; sTu, Shimizu, and Berg 2008)*.

R_a_ facilitates a self-enhancing component, “S”, which represents the VGCC and TRPV channels combined. Their dynamics can be described by:

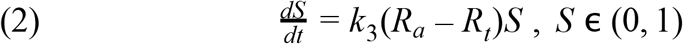

Thus, when R_a_ crosses a critical activation threshold *R*_*t*_ (0 < *R*_*t*_ < 1), S will grow/fall exponentially. S is approximated to form a self-enhancing motif where a small calcium influx generated by the Ra-mediated opening of TRPV channels initiates further opening and self-amplification of VGCCs, through which most of the calcium enters the cell(Larsch et al. 2015; Liu et al. 2018).

S values are limited by code to the interval *S* ∊ (0, 1], to simplify a more realistic sigmoidal behaviour. Its lower limit is a small (<<1) positive number and its exact limit does not affect model performance (not shown). Thus, S produces a “two-state” switch where the neuron is either active or inactive. For this, the time constant 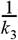, needs to be smaller than the other time constants in the system, though its specific values do not significantly affect the model performance.

In equation 3, S facilitates calcium accumulation in the cell:

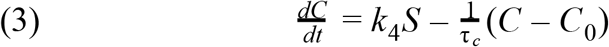

Calcium enters through open channels (S) and is removed by calcium pumps. The removal time constant, τ _*c*_ was approximated experimentally to be in the order of several seconds (Itskovits, Ruach, and Zaslaver 2018). *C*_0_ is the calcium level at rest so that *C*≥*C*_0_. The calcium in the cell facilitates the Inhibition term, according to equation 4:

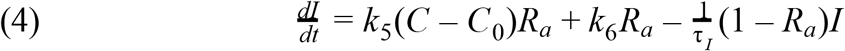

Where *k*_5_, *k*_6_ and τ _*I*_ are positive constants. Two components increase inhibition levels: the first depends on calcium (with the constant *k*_5_) and the second is calcium independent (denoted with the constant *k*_6_). The calcium-dependent term is consistent with homologous adaptation by G protein-coupled receptor kinases (GRKs) that phosphorylate only active receptors (which is why the term is multiplied by *R*_*a*_). The second term depends on receptor activity only and is consistent with phosphorylation driven by second messenger-stimulated kinases(Lefkowitz 1998).

These two terms fulfill two different features in the system. To achieve robust exact adaptation, the circuit’s output needs to directly inhibit the receptor(Yi et al. 2000). Therefore, as calcium directly correlates with the synaptic output(Katz and Miledi 1970; Llinás, Steinberg, and Walton 1976), its regulation of receptor inhibition underlies robust exact adaptation. However, as calcium levels are transient, adaptation should also depend on the degree of receptor activation to promote inhibition for as long as the stimulus is present. Our experimental results support this logic as we find that calcium levels correlate with adaptation magnitude, and this adaptation persists even after calcium removal (**Figure 4**). These results suggest two separate adaptation mechanisms: one that is calcium dependent and the other which relies on the receptor’s activity.

The last term in equation 4 denotes inhibition removal which depends on inactive receptors only (thus multiplying ‘I’ by (1-*R*_*a*_)), similar to methylation dynamics in *E. coli(Shimizu, Tu, and Berg 2010)*. To close the negative feedback loop, this inhibition (I) regulates receptor activation according to equation 1.

### Mathematical analysis of the model’s steady state shows how exact adaptation is achieved

The mathematical formulation of exact adaptation requires that following a step increase in the attractant, the steady state value of the circuit output must return to their basal pre-stimulation levels. To examine this transition we set *L* → *L*_1_ and require 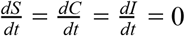. From equation 2, a stable equilibrium can only be reached by *S* = 1 or *S* → 0. A steady state in which *S* = 1 represents a non-pulsatile response, in which calcium stays at its maximum, and can be obtained by choices of parameters where calcium-mediated inhibition is low (as in *tax-6* mutants, **Supplementary figure 6**). A steady state in which *S* → 0 may lead to exact adaptation as follows: by plugging *S* = 0 in equation 3, we get a steady-state calcium level that does not depend on the input: *C*_*SS*_ = *C*_0_. Furthermore, plugging these results in equation 4, yields:

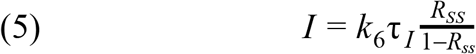

If we plug *I* in equation 1, we get a closed form for the receptor activation during steady state:

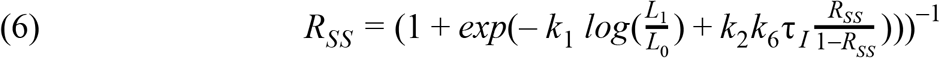

This relation can be solved numerically, and by plugging this solution back to equation 5, we get the steady state inhibition level, *I*_*SS*_. We can see that steady-state activation and inhibition do depend on the input level L_1_, thus serving as a ‘memory’ of the input, even after output returns to basal levels.

Considering dynamics, when responding to a step-function input, receptor activation will rise above *R*_*t*_, elicit an output pulse that will, in turn, increase inhibition. Inhibition will eventually decrease receptor activation to terminate the pulse. In the regime where *R*_*s*s_ > *R*_*t*_, receptor activation will rise and cross *R*_*t*_ before reaching steady state, and thus elicit another pulse. Therefore, a strict requirement for exact adaptation in this simple model is *R*_*ss*_ < *R*_*t*_.

### How smooth gradients of input are encoded as a pulsatile response that adapts to the gradients first derivative?

Our model translates smooth gradients of the input into a pulsatile response output: Increased ligand levels lead to receptor activation (eq. 1). At this stage, since there is still no output, inhibition is relatively low (eq. 4), and for simplification, we assume its effect on receptor activation to be negligible. When receptor activation reaches its threshold level, *R*_*t*_, a pulse of calcium is generated (eqs. 2 and 3). High levels of calcium rapidly increase inhibition (eq. 4), though at this time scale, ligand concentrations remain relatively constant. The rapid increase in inhibition causes a sharp decline in receptor activation levels (eq. 1), and when decreased below *R*_*t*_, the pulse terminates (eq. 2,3). Calcium is being removed and hence calcium-mediated inhibition slowly decreases (eq. 4), while ligand concentrations increase to elicit another pulse (as can be seen in **Figure 2b**).

To explain how the above pulsatile response adapts to the first derivative of the input gradient, we analyze the model with the following assumptions: (1) a calcium pulse is an immediate event, in which ligand levels remain constant, and the inhibition rises to a new higher level; (2) the change in inhibition during a pulse always results in a constant decrease of receptor activation; (3) between pulses, inhibition remains constant, so that only the ligand can change receptor activation.

We assume a first pulse occurs when activation reaches *R*_*t*_ at a ligand level *L*_1_ and inhibition level *I*_1_. This pulse increases inhibition to a new *I*_2_ level, so that receptor activation falls to a new level *R*_0_ < *R*_*t*_:

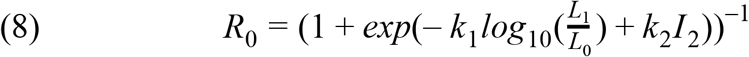

To elicit an additional pulse, the ligand concentration has to rise to a new level, *L*_2_, so that receptor activation reaches *R*_*t*_ once again. Inhibition is assumed to stay constant, thus when a second pulse is produced:

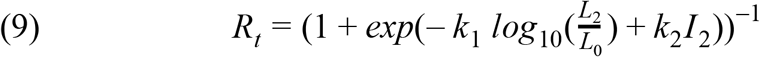

Isolating *I*_2_ in equation 8, and placing in equation 9 yields:

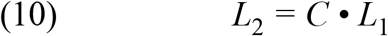

Where 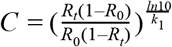 is greater than 1 since *R*_*t*_ > *R*_0_ and *k*_1_ > 0. Thus to produce the consecutive pulse, ligand levels need to increase by a factor *C*. For a linear increase in ligand levels, *L* = α*t*; this means that the n’th pulse time, and the time between consecutive pulses will be:

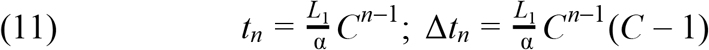

Thus, the time interval between consecutive pulses grows with the number of pulses. The same consideration will also dictate less pulses in the second half of a sigmoidal gradient (decreasing first derivatives of the gradient), effectively exhibiting adaptation to the first derivative.

### Integrating the model with known dynamics of voltage gated ion channels in AWA

To accurately account for the main voltage-gated ion channels in the AWA neuron, we integrated our model with a model developed in the Bargmann lab based on intracellular recordings of the AWA neuron(Liu et al. 2018). This model takes electrical current influx as an input, and simulates the dynamics of ion channels and membrane potential in the cell. To integrate this model into our simulations, the electrical current influx was considered to be proportional to our self-enhancing motif:

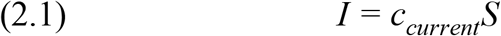

Where *I* is the current influx, *S* is the self-enhancing motif, and *c*_*current*_ is the proportion constant which determines the connectivity between the models. Thus, the current influx is provided as the input to the model:

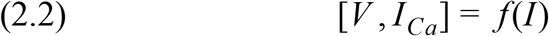

Where *f* is the model, *V* is the membrane potential and *I*_*Ca*_ is the calcium influx entering the cell. Calcium influx is therefore plugged into equation 3, instead of *S*:

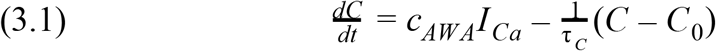

Where *c*_*AWA*_ is a constant that changes the electrical current units to concentration changes while accounting for the volume of the AWA neuron. To determine this constant, we approximated the cell volume based on our microscopy images to be: *V*_*AWA*_ = 9 • 4 • 4 μ*m*^3^ ~ 2 • 10^−13^ *L*, and used the following transformation:

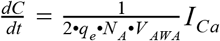

Where *q*_*e*_ is the elementary charge, *N*_*A*_ is the Avogadro number, and 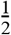 accounts for the charge of a calcium ion. Overall, in the detailed integrated model, equations 2.1 and 2.2 replace equation 2 (in the original model), and equation 3.1 replaces equation 3 (in the original model).

## References

Alon, U., Surette, M. G., Barkai, N., & Leibler, S. (1999). Robustness in bacterial chemotaxis. In Nature (Vol. 397, Issue 6715, pp. 168–171). https://doi.org/10.1038/16483

Bargmann, C. (2006). Chemosensation in C. elegans. In WormBook. https://doi.org/10.1895/wormbook.1.123.1

Bargmann, C. I., Hartwieg, E., & Robert Horvitz, H. (1993). Odorant-selective genes and neurons mediate olfaction in C. elegans. In Cell (Vol. 74, Issue 3, pp. 515–527). https://doi.org/10.1016/0092-8674(93)80053-h

Barkai, N., & Leibler, S. (1997). Robustness in simple biochemical networks. Nature, 387(6636), 913–917.

Berg, H. C., & Tedesco, P. M. (1975). Transient response to chemotactic stimuli in Escherichia coli. Proceedings of the National Academy of Sciences of the United States of America, 72(8), 3235–3239.

Block, S. M., Segall, J. E., & Berg, H. C. (1983). Adaptation kinetics in bacterial chemotaxis. In Journal of Bacteriology (Vol. 154, Issue 1, pp. 312–323). https://doi.org/10.1128/jb.154.1.312-323.1983

Choi, J. I., Yoon, K.-H., Kalichamy, S. S., Yoon, S.-S., & Lee, J. I. (2016). A natural odor attraction between lactic acid bacteria and the nematode Caenorhabditis elegans. In The ISME Journal (Vol. 10, Issue 3, pp. 558–567). https://doi.org/10.1038/ismej.2015.134

Chronis, N., Zimmer, M., & Bargmann, C. I. (2007). Microfluidics for in vivo imaging of neuronal and behavioral activity in Caenorhabditis elegans. Nature Methods, 4(9), 727–731.

Colbert, H. A., & Bargmann, C. I. (1995). Odorant-specific adaptation pathways generate olfactory plasticity in C. elegans. In Neuron (Vol. 14, Issue 4, pp. 803–812). https://doi.org/10.1016/0896-6273(95)90224-4

Cook, S. J., Jarrell, T. A., Brittin, C. A., Wang, Y., Bloniarz, A. E., Yakovlev, M. A., Nguyen, K. C. Q., Tang, L. T.-H., Bayer, E. A., Duerr, J. S., Bülow, H. E., Hobert, O., Hall, D. H., & Emmons, S. W. (2019). Whole-animal connectomes of both Caenorhabditis elegans sexes. Nature, 571(7763), 63–71.

Edelstein, A. D., Tsuchida, M. A., Amodaj, N., Pinkard, H., Vale, R. D., & Stuurman, N. (2014). Advanced methods of microscope control using μManager software. Journal of Biological Methods, 1(2). https://doi.org/10.14440/jbm.2014.36

Fechner, G. T. (n.d.). Elements of psychophysics, 1860. In Readings in the history of psychology. (pp. 206–213). https://doi.org/10.1037/11304-026

Ferrell, J. E., Jr. (2016). Perfect and Near-Perfect Adaptation in Cell Signaling. Cell Systems, 2(2), 62–67.

Ferrell, J. E., Jr, & Machleder, E. M. (1998). The biochemical basis of an all-or-none cell fate switch in Xenopus oocytes. Science, 280(5365), 895–898.

Fukuto, H. S., Ferkey, D. M., Apicella, A. J., Lans, H., Sharmeen, T., Chen, W., Lefkowitz, R. J., Jansen, G., Schafer, W. R., & Hart, A. C. (2004). G Protein-Coupled Receptor Kinase Function Is Essential for Chemosensation in C. elegans. In Neuron (Vol. 42, Issue 4, pp. 581–593). https://doi.org/10.1016/s0896-6273(04)00252-1

Gaudry, Q., Nagel, K. I., & Wilson, R. I. (2012). Smelling on the fly: sensory cues and strategies for olfactory navigation in Drosophila. Current Opinion in Neurobiology, 22(2), 216–222.

Gordus, A., Pokala, N., Levy, S., Flavell, S. W., & Bargmann, C. I. (2015). Feedback from network states generates variability in a probabilistic olfactory circuit. Cell, 161(2), 215–227.

Hart, A., & Chao, M. (2009). From Odors to Behaviors in Caenorhabditis Elegans. In Frontiers in Neuroscience (pp. 1–33). https://doi.org/10.1201/9781420071993-c1

Hobert, O. (2013). The neuronal genome of Caenorhabditis elegans. In WormBook (pp. 1–106). https://doi.org/10.1895/wormbook.1.161.1

Itskovits, E., Ruach, R., & Zaslaver, A. (2018). Concerted pulsatile and graded neural dynamics enables efficient chemotaxis in C. elegans. Nature Communications, 9(1), 2866.

Kahn-Kirby, A. H., Dantzker, J. L. M., Apicella, A. J., Schafer, W. R., Browse, J., Bargmann, C. I., & Watts, J. L. (2004). Specific polyunsaturated fatty acids drive TRPV-dependent sensory signaling in vivo. Cell, 119(6), 889–900.

Katz, B., & Miledi, R. (1970). Further study of the role of calcium in synaptic transmission. In The Journal of Physiology (Vol. 207, Issue 3, pp. 789–801). https://doi.org/10.1113/jphysiol.1970.sp009095

Kuhara, A., Inada, H., Katsura, I., & Mori, I. (2002). Negative regulation and gain control of sensory neurons by the C. elegans calcineurin TAX-6. Neuron, 33(5), 751–763.

Larsch, J., Flavell, S. W., Liu, Q., Gordus, A., Albrecht, D. R., & Bargmann, C. I. (2015). A Circuit for Gradient Climbing in C. elegans Chemotaxis. Cell Reports, 12(11), 1748–1760.

Larsch, J., Ventimiglia, D., Bargmann, C. I., & Albrecht, D. R. (2013). High-throughput imaging of neuronal activity in Caenorhabditis elegans. Proceedings of the National Academy of Sciences of the United States of America, 110(45), E4266–E4273.

Laughlin, S. B. (1989). The role of sensory adaptation in the retina. The Journal of Experimental Biology. https://jeb.biologists.org/content/146/1/39.short

Lazova, M. D., Ahmed, T., Bellomo, D., Stocker, R., & Shimizu, T. S. (2011). Response rescaling in bacterial chemotaxis. Proceedings of the National Academy of Sciences of the United States of America, 108(33), 13870–13875.

Lefkowitz, R. J. (1998). G Protein-coupled Receptors. In Journal of Biological Chemistry (Vol. 273, Issue 30, pp. 18677–18680). https://doi.org/10.1074/jbc.273.30.18677

Levy, S., & Bargmann, C. I. (2020). An Adaptive-Threshold Mechanism for Odor Sensation and Animal Navigation. Neuron, 105(3), 534–548.e13.

Liu, Q., Kidd, P. B., Dobosiewicz, M., & Bargmann, C. I. (2018). C. elegans AWA Olfactory Neurons Fire Calcium-Mediated All-or-None Action Potentials. In Cell (Vol. 175, Issue 1, pp. 57–70.e17). https://doi.org/10.1016/j.cell.2018.08.018

Llinás, R., Steinberg, I. Z., & Walton, K. (1976). Presynaptic calcium currents and their relation to synaptic transmission: voltage clamp study in squid giant synapse and theoretical model for the calcium gate. Proceedings of the National Academy of Sciences of the United States of America, 73(8), 2918–2922.

Luo, L., Wen, Q., Ren, J., Hendricks, M., Gershow, M., Qin, Y., Greenwood, J., Soucy, E. R., Klein, M., Smith-Parker, H. K., Calvo, A. C., Colón-Ramos, D. A., Samuel, A. D. T., & Zhang, Y. (2014). Dynamic Encoding of Perception, Memory, and Movement in a C. elegans Chemotaxis Circuit. In Neuron (Vol. 82, Issue 5, pp. 1115–1128). https://doi.org/10.1016/j.neuron.2014.05.010

Marder, E., & Goaillard, J.-M. (2006). Variability, compensation and homeostasis in neuron and network function. Nature Reviews. Neuroscience, 7(7), 563–574.

Murlis, J., Elkinton, J. S., & Cardé, R. T. (1992). Odor Plumes and How Insects Use Them. In Annual Review of Entomology (Vol. 37, Issue 1, pp. 505–532). https://doi.org/10.1146/annurev.en.37.010192.002445

Olsman, N., & Goentoro, L. (2016). Allosteric proteins as logarithmic sensors. Proceedings of the National Academy of Sciences of the United States of America, 113(30), E4423–E4430.

Pierce-Shimomura, J. T., Morse, T. M., & Lockery, S. R. (1999). The fundamental role of pirouettes in Caenorhabditis elegans chemotaxis. The Journal of Neuroscience: The Official Journal of the Society for Neuroscience, 19(21), 9557–9569.

Pritz, C. O., Itskovits, E., Bokman, E., Ruach, R., Gritsenko, V., Nelken, T., Menasherof, M., Azulay, A., & Zaslaver, A. (n.d.). Principles for coding associative memories in a compact neural network. https://doi.org/10.1101/2020.06.20.162818

Rahi, S. J., Larsch, J., Pecani, K., Katsov, A. Y., Mansouri, N., Tsaneva-Atanasova, K., Sontag, E. D., & Cross, F. R. (2017). Oscillatory stimuli differentiate adapting circuit topologies. Nature Methods, 14(10), 1010–1016.

Sengupta, P., Chou, J. H., & Bargmann, C. I. (1996). odr-10 encodes a seven transmembrane domain olfactory receptor required for responses to the odorant diacetyl. Cell, 84(6), 899–909.

Shimizu, T. S., Tu, Y., & Berg, H. C. (2010). A modular gradient-sensing network for chemotaxis in Escherichia coli revealed by responses to time-varying stimuli. Molecular Systems Biology, 6, 382.

Shirley, S. G., Polak, E. H., Edwards, D. A., Wood, M. A., & Dodd, G. H. (1987). The effect of concanavalin A on the rat electro-olfactogram at various odorant concentrations. Biochemical Journal, 245(1), 185–189.

Sourjik, V., & Wingreen, N. S. (2012). Responding to chemical gradients: bacterial chemotaxis. Current Opinion in Cell Biology, 24(2), 262–268.

Stern, S., Kirst, C., & Bargmann, C. I. (2017). Neuromodulatory Control of Long-Term Behavioral Patterns and Individuality across Development. Cell, 171(7), 1649–1662.e10.

Sulston, J. E., & Brenner, S. (1974). The DNA of Caenorhabditis elegans. Genetics, 77(1), 95–104.

Tu, Y. (2013). Quantitative modeling of bacterial chemotaxis: signal amplification and accurate adaptation. Annual Review of Biophysics, 42, 337–359.

Tu, Y., & Rappel, W.-J. (2018). Adaptation of Living Systems. Annual Review of Condensed Matter Physics, 9, 183–205.

Tu, Y., Shimizu, T. S., & Berg, H. C. (2008). Modeling the chemotactic response of Escherichia coli to time-varying stimuli. Proceedings of the National Academy of Sciences of the United States of America, 105(39), 14855–14860.

van As, W., Kauer, J. S., Menco, B. P. M., & Köster, E. P. (1985). Quantitative aspects of the electro-olfactogram in the tiger salamander. In Chemical Senses (Vol. 10, Issue 1, pp. 1–21). https://doi.org/10.1093/chemse/10.1.1-b

Vinograd, A., Livneh, Y., & Mizrahi, A. (2017). History-Dependent Odor Processing in the Mouse Olfactory Bulb. The Journal of Neuroscience: The Official Journal of the Society for Neuroscience, 37(49), 12018–12030.

Vries, L. D., De Vries, L., Zheng, B., Fischer, T., Elenko, E., & Farquhar, M. G. (2000). The Regulator of G Protein Signaling Family. In Annual Review of Pharmacology and Toxicology (Vol. 40, Issue 1, pp. 235–271). https://doi.org/10.1146/annurev.pharmtox.40.1.235

White, J. G., Southgate, E., Thomson, J. N., & Brenner, S. (1986). The structure of the nervous system of the nematode Caenorhabditis elegans. In Philosophical Transactions of the Royal Society of London. B, Biological Sciences (Vol. 314, Issue 1165, pp. 1–340). https://doi.org/10.1098/rstb.1986.0056

Zhang, X.-P., Cheng, Z., Liu, F., & Wang, W. (2007). Linking fast and slow positive feedback loops creates an optimal bistable switch in cell signaling. Physical Review. E, Statistical, Nonlinear, and Soft Matter Physics, 76(3 Pt 1), 031924.

Zufall, F. (2000). The Cellular and Molecular Basis of Odor Adaptation. In Chemical Senses (Vol. 25, Issue 4, pp. 473–481). https://doi.org/10.1093/chemse/25.4.473

## References

Itskovits, Eyal, Rotem Ruach, and Alon Zaslaver. 2018. “Concerted Pulsatile and Graded Neural Dynamics Enables Efficient Chemotaxis in C. Elegans.” Nature Communications. https://doi.org/10.1038/s41467-018-05151-2.

Katz, B., and R. Miledi. 1970. “Further Study of the Role of Calcium in Synaptic Transmission.” The Journal of Physiology. https://doi.org/10.1113/jphysiol.1970.sp009095.

Larsch, Johannes, Steven W. Flavell, Qiang Liu, Andrew Gordus, Dirk R. Albrecht, and Cornelia Bargmann. 2015. “A Circuit for Gradient Climbing in C. Elegans Chemotaxis.” Cell Reports 12 (11): 1748–60.

Lefkowitz, R. J. 1998. “G Protein-Coupled Receptors. III. New Roles for Receptor Kinases and Beta-Arrestins in Receptor Signaling and Desensitization.” The Journal of Biological Chemistry 273 (30): 18677–80.

Liu, Qiang, Philip B. Kidd, May Dobosiewicz, and Cornelia I. Bargmann. 2018. “C. Elegans AWA Olfactory Neurons Fire Calcium-Mediated All-or-None Action Potentials.” Cell. https://doi.org/10.1016/j.cell.2018.08.018.

Llinás, R., I. Z. Steinberg, and K. Walton. 1976. “Presynaptic Calcium Currents and Their Relation to Synaptic Transmission: Voltage Clamp Study in Squid Giant Synapse and Theoretical Model for the Calcium Gate.” Proceedings of the National Academy of Sciences of the United States of America 73 (8): 2918–22.

Rahi, Sahand Jamal, Johannes Larsch, Kresti Pecani, Alexander Y. Katsov, Nahal Mansouri, Krasimira Tsaneva-Atanasova, Eduardo D. Sontag, and Frederick R. Cross. 2017. “Oscillatory Stimuli Differentiate Adapting Circuit Topologies.” Nature Methods 14 (10): 1010–16.

Shimizu, Thomas S., Yuhai Tu, and Howard C. Berg. 2010. “A Modular Gradient-Sensing Network for Chemotaxis in Escherichia Coli Revealed by Responses to Time-Varying Stimuli.” Molecular Systems Biology 6 (June): 382.

Tu, Yuhai, Thomas S. Shimizu, and Howard C. Berg. 2008. “Modeling the Chemotactic Response of Escherichia Coli to Time-Varying Stimuli.” Proceedings of the National Academy of Sciences of the United States of America 105 (39): 14855–60.

Yi, T. M., Y. Huang, M. I. Simon, and J. Doyle. 2000. “Robust Perfect Adaptation in Bacterial Chemotaxis through Integral Feedback Control.” Proceedings of the National Academy of Sciences of the United States of America 97 (9): 4649–53.

